# The *C. difficile clnRAB* operon initiates adaptations to the host environment in response to LL-37

**DOI:** 10.1101/347286

**Authors:** Emily C. Woods, Adrianne N. Edwards, Shonna M. McBride

## Abstract

To cause disease, *Clostridioides* (*Clostridium*) *difficile* must resist killing by innate immune effectors in the intestine, including the host antimicrobial peptide, cathelicidin (LL-37). The mechanisms that enable *C. difficile* to adapt to the intestine in the presence of antimicrobial peptides are unknown. Expression analyses revealed an operon, *CD630_16170-CD630_16190* (*clnRAB*), which is highly induced by LL-37 and is not expressed in response to other cell-surface active antimicrobials. This operon encodes a predicted transcriptional regulator (*clnR*) and an ABC transporter system (*clnAB*), all of which are required for function. Analyses of a *clnR* mutant indicate that ClnR is a pleiotropic regulator that directly binds to LL-37 and controls expression of numerous genes, including many involved in metabolism, cellular transport, signaling, gene regulation, and pathogenesis. The data suggest that ClnRAB is a novel regulatory mechanism that senses LL-37 as a host signal and regulates gene expression to adapt to the host intestinal environment during infection.

**Author Summary:** *C. difficile* is a major nosocomial pathogen that causes severe diarrheal disease. Though *C. difficile* is known to inhabit the human gastrointestinal tract, the mechanisms that allow this pathogen to adapt to the intestine and survive host defenses are not known. In this work, we investigated the response of *C. difficile* to the host defense peptide, LL-37, to determine the mechanisms underlying host adaptation and survival. Expression analyses revealed a previously unknown locus, which we named *clnRAB*, that is highly induced by LL-37 and acts as a global regulator of gene expression in *C. difficile*. Mutant analyses indicate that ClnRAB is a novel regulatory system that senses LL-37 as a host signal to regulate adaptation to the intestinal environment.

## INTRODUCTION

*Clostridioides difficile* (formerly *Clostridium difficile*) poses a serious, ongoing, public health threat. *C. difficile* infection (CDI) results in mild to severe diarrhea and leads to approximately 29,000 deaths each year in the United States (1). Patients are typically infected after treatment with antibiotics, which disrupt the intestinal microbiota that provide colonization resistance against CDI by competition and release of antimicrobial peptides (AMPs) (1, 2).

The host innate immune system also plays an important role in the prevention of infections. A critical feature of this defense is the production of AMPs, including defensins, cathelicidin (LL-37), and lysozyme (3, 4). LL-37, a cationic AMP, is of particular importance in CDI because it is not only produced constitutively in the colon by the colonic epithelium, but is also released in high levels from neutrophils, which are a key component of the initial immune response to CDI (5). LL-37 is stored in neutrophil granules at a concentration of ∼6 µg/ml and has been reported to reach levels of 15 µg/ml in the lungs of cystic fibrosis patients and ∼5 µg per gram of feces during shigellosis (6-8). LL-37 forms an amphipathic alpha-helical structure that can insert into bacterial membranes and cause bacterial cell death (9-11). Common bacterial resistance mechanisms to LL-37 include cell surface modifications that prevent LL-37 access to the bacterial surface, efflux pumps that eliminate LL-37 that enters the bacteria, secreted proteases that degrade LL-37, and modulation of host production of LL-37 (5).

*C. difficile* demonstrates inducible resistance to LL-37, and current epidemic ribotypes have higher levels of resistance to LL-37 than other ribotypes (12). Although *C. difficile* resistance to LL-37 is documented, no clear homologs of known resistance mechanisms are apparent in the genome and no additional LL-37 resistance mechanisms have been identified. Moreover, the mechanisms by which *C. difficile* responds to this and other innate immune factors are poorly understood. We hypothesized that *C. difficile* recognizes and responds to LL-37 and that this response occurs, at least in part, at the level of transcription.

In this study, we determined the transcriptional response of *C. difficile* to the host peptide, LL-37. We identified an operon, *CD630_16170-CD630_16190* (herein named *clnRAB*), that was highly induced by LL-37 and was not expressed in response to other cell-surface active antimicrobials. This operon encodes a predicted GntR-family transcriptional regulator (*clnR*) and an ABC transporter system (*clnAB*). We determined that the ClnR regulator represses *clnRAB* expression and is also necessary for LL-37 dependent induction of *clnRAB* transcription. Transcriptional analyses of a *clnR* mutant indicated that ClnR is a global regulator that controls the expression of numerous genes including toxins, alternative metabolism pathways, transporters, and transcriptional regulators. Growth analyses revealed that exposure to LL-37 modifies the metabolism of *C. difficile* and that this response occurs through ClnR. In addition, we observed that both a *clnR* and a *clnAB* mutant are more virulent in the hamster model of infection and a *clnR* mutant is defective at colonization in the mouse model of infection. *In vitro* analyses confirmed that ClnR is a DNA-binding transcriptional regulator that directly controls expression of the *cln* operon and other ClnR-regulated genes. Further, we observed specific binding of ClnR to LL-37, verifying the direct interaction of these factors. Based on these data, we propose that LL-37 acts as a host signal that is transmitted through ClnRAB, enabling *C. difficile* to regulate global gene expression to adapt to the intestinal environment during infection.

## RESULTS

### Discovery of an operon that is highly induced by LL-37

To test the hypothesis that *C. difficile* responds to LL-37 through changes in gene transcription, we performed RNA-seq analysis on bacteria grown with or without sub-MIC levels of LL-37 to determine which genes were differentially expressed. RNA-seq analysis revealed 228 genes that were differentially expressed at least 2-fold and with *p* < 0.05, including 107 genes that were induced and 121 genes that were down-regulated in the presence of LL-37 (**Table S1**). Genes differentially expressed in LL-37 include loci predicted to encode metabolic pathway components, nutrient acquisition mechanisms, transcriptional regulators, multidrug transporters, antibiotic resistance factors, conjugation-associated proteins, and genes of unknown function. Notably, several of these loci were previously investigated in *C. difficile* and found to contribute to growth, antimicrobial resistance or virulence, including genes involved in succinate, glucose, fructose, mannitol, ethanolamine, butyrate, acetyl-CoA and amino acid metabolism, oligopeptide permeases, elongation factor (EF-G), ferredoxin oxidoreductase, and ECF sigma factors (13-23).

Of the differentially regulated genes identified, the most highly induced by LL-37 were three genes comprising an apparent operon: *CD630_16170-16190*. These genes encode a putative GntR-family transcriptional regulator (*CD630_16170, clnR*) and a downstream ABC transporter system composed of an ATP-binding component (*CD630_16180, clnA*) and a permease (*CD630_16190, clnB*). The results of the RNA-seq analyses for *clnRAB* expression were verified by qRT-PCR in the 630Δ*erm* strain and for the epidemic 027 ribotype strain, R20291 (**Fig. 1**). Based on the substantial induction of these genes and their resemblance to antimicrobial response systems, we pursued the function of this operon further. Transcriptional analysis of strains grown in increasing concentrations of LL-37 demonstrated that expression of each of these genes increased in a dose-dependent manner for both strains, illustrating that these genes are similarly expressed and regulated in diverse *C. difficile* isolates (**Fig. 1**). Using nested PCR from *C. difficile* cDNA templates, we also confirmed that the *CD630_16170-16190* genes are transcribed as an operon (**Fig. S1**).

**Figure 1.**
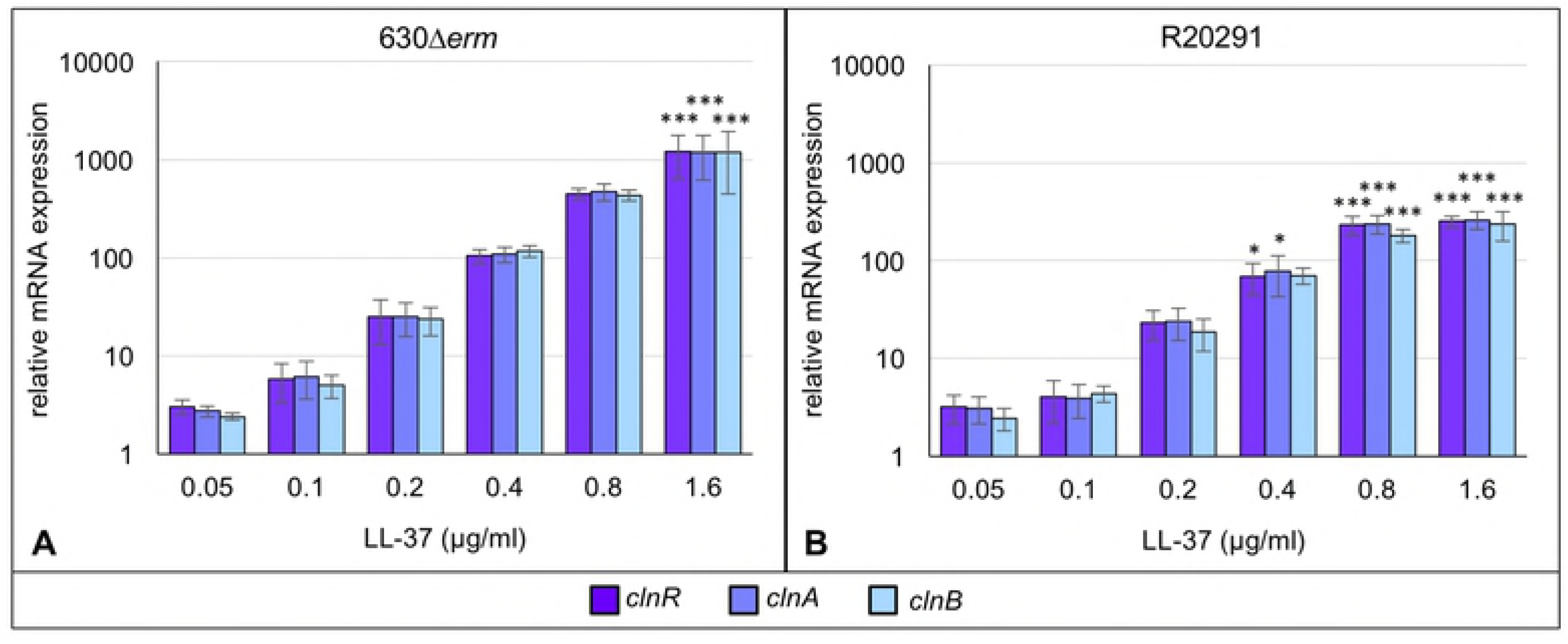
Expression of *CD630_16170-CD630_16190* is induced by LL-37 in a dose-dependent manner. Active cultures of A) 630Δ*erm* and B) R20291 were grown in BHIS or BHIS with 0.05, 0.1, 0.2, 0.4, 0.8 or 1.6 µg/ml LL-37. Samples were harvested, cDNA generated, and qRT-PCR performed as described in Methods. mRNA levels are normalized to expression levels in BHIS alone. Bars represent the mean and standard deviation for at least three biological replicates. Expression levels of each gene were analyzed by one-way ANOVA and Dunnett’s multiple comparisons test, comparing to expression without LL-37. Adjusted *p* values indicated by * ≤ 0.05, ** ≤ 0.01, *** ≤ 0.001.

### The *cln* operon does not contribute significantly to LL-37 resistance

Given the high level of induction of the *clnRAB* operon in LL-37, and because transporters are common antimicrobial resistance mechanisms, we hypothesized that ClnAB may confer resistance to LL-37. To test this, we generated insertional disruptions in the *CD630_16170* (*clnR::erm*; herein *clnR* null mutant) and *CD630_16180* (*clnA::erm*, herein *clnAB* null mutant) coding sequences (**Fig. S2A**), and analyzed the ability of the mutants to grow in the presence of LL-37 (**Fig. 2**). Although the *clnR* mutant has a minor growth defect when grown in BHIS alone, both the *clnR* and *clnA* mutants grew slightly better than the parent strain in 2.5 µg/ml LL-37. Evaluation of the minimum inhibitory concentrations (MICs) and minimum bactericidal concentrations (MBCs) for LL-37 in these strains revealed no observable differences in either the MIC or MBC for the *clnR* or *clnAB* null mutant in comparison to the parent strain (**Table S2**). Considering that some antimicrobial transporter mechanisms are activated by and confer resistance to multiple classes of antimicrobials, we investigated the resistance of both mutants to other cell-surface acting compounds. Neither the *clnR* nor the *clnAB* mutant had altered MIC values for any other cell surface-active antimicrobial tested (**Table S3**). These findings indicate that the *cln* operon does not play a significant role in resistance to LL-37 or other tested cell-surface active antimicrobials.

**Figure 2.**
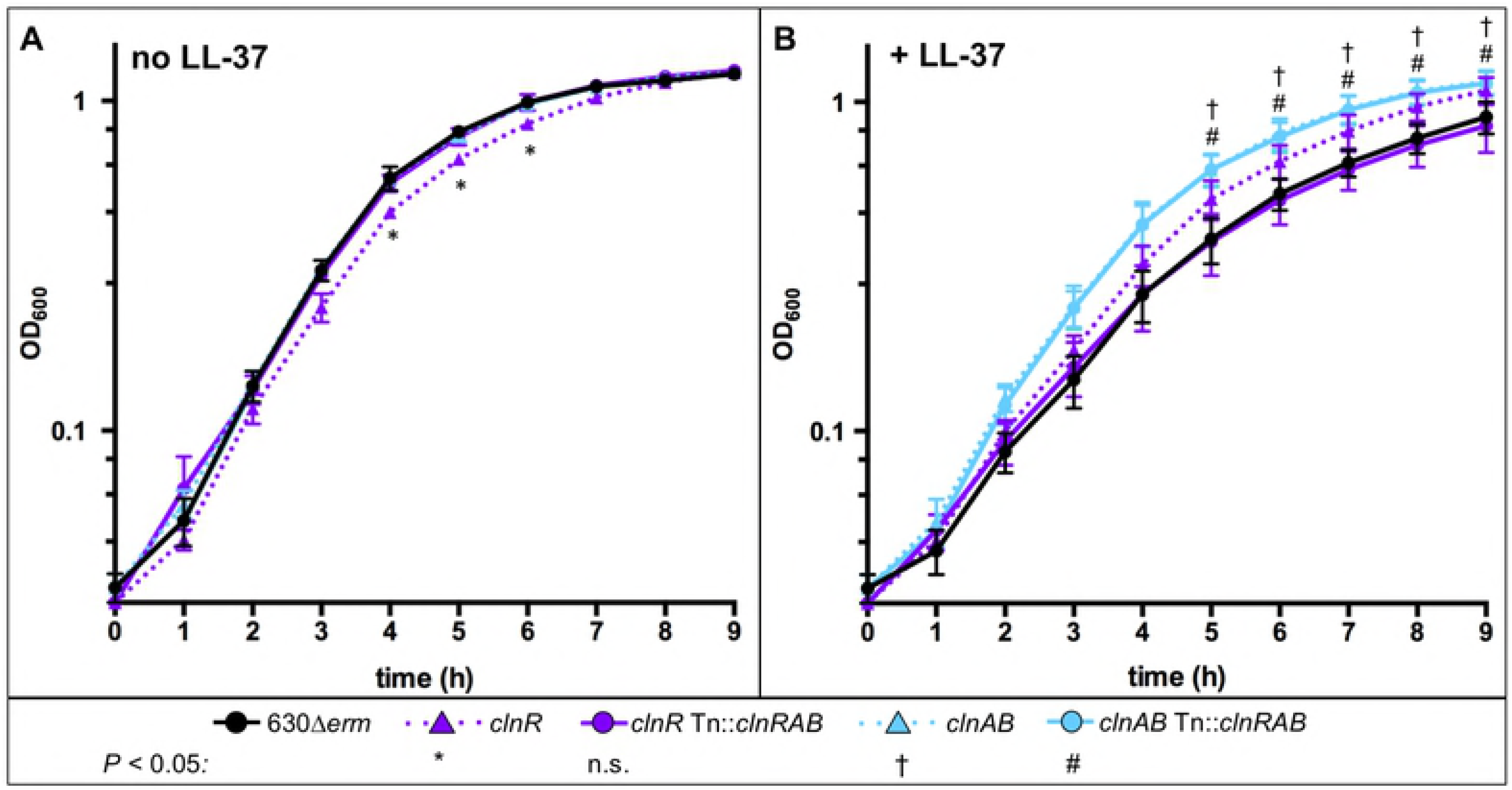
Growth of *clnR* and *clnAB* mutants with and without LL-37. Active cultures of 630Δ*erm* (black circles), *clnR* (MC885; purple triangles), *clnR* Tn::*clnRAB* (MC950; purple circles), *clnAB* (MC935; blue triangles), *clnAB* Tn::*clnRAB* (MC953; blue circles) were diluted to an OD_600_ of 0.05 in **A)** BHIS alone or **B)** BHIS with 2.5 µg/ml LL-37. Graphs represent the means +/− SEM from three independent replicates. Data were analyzed by two-way ANOVA with Dunnett’s multiple comparisons test, comparing to 630Δ*erm* at the same time point. * † # indicate adjusted *p* < 0.05 for individual strains, as indicated; ns: not significant.

### Induction of *clnRAB* is specific to LL-37-like cathelicidins

The induction of the *cln* operon suggested that this locus was responsive to LL-37; however, antimicrobials may induce changes in bacterial gene expression as a general stress response or due to disruptions in cellular processes (24, 25). To determine whether the induction of this operon is specific to LL-37 or a general response to cellular stress, we evaluated the expression of *clnR* in the presence of a variety of other antimicrobial compounds (**Table 1**). Transcription of *clnR* was also induced when *C. difficile* was exposed to the mouse cathelicidin, mCRAMP, but *clnR* was not induced in the presence of sequence-scrambled LL-37 or the sheep cathelicidin, SMAP-29, which is less similar to LL-37 than mCRAMP (**Fig. S3**) (26-28). Similarly, none of the other cell-surface-active antimicrobials tested (lysozyme, ampicillin, vancomycin, nisin, or polymyxin B) induced *clnR* expression (**Table 1**). These results indicate that induction of the *clnRAB* operon is dependent on the specific sequence of LL-37 and is not caused by antimicrobial-induced cell-surface stress. Accordingly, we named this operon *clnRAB* to reflect the specificity of the induction in response to LL-37 and similar **<u>C</u>**athe**<u>l</u>**icidi**<u>n</u>**s.

**Table 1.** Induction of *clnR* is specific to LL-37-like cathelicidins

| Antimicrobial<sup>a</sup> | Relative *clnR* expression<sup>b, c</sup> |
| --- | --- |
| LL-37 | **530.1 ± 110.5** |
| scrambled LL-37 | 1.7 ± 0.8 |
| mCRAMP | **111.4 ± 27.3** |
| SMAP-29 | 0.7 ± 0.1 |
| Lysozyme | 1.1 ± 0.3 |
| Ampicillin | 0.9 ± 0.2 |
| Vancomycin | 0.9 ± 0.1 |
| Nisin | 1.1 ± 0.4 |
| Polymyxin B | 1.2 ± 0.2 |

<sup>a</sup>Concentrations of antimicrobials used: 2 µg/ml LL-37, 2 µg/ml scrambled LL-37, 2 µg/ml mCRAMP, 0.35 µg/ml SMAP-29, 1 mg/ml lysozyme, 4 µg/ml ampicillin, 0.5 µg/ml vancomycin, 7.5 µg/ml nisin, 200 µg/ml polymyxin B
<sup>b</sup>Relative expression determined by qRT-PCR and normalized to 630Δ*erm* grown in BHIS without antimicrobials. Values are the mean of three replicates ± standard error of the mean.
<sup>c</sup>Bolded values indicate significant difference (adjusted *P* value < 0.05) from 630Δ*erm* grown in BHIS without antimicrobials and analyzed by one-way ANOVA and Dunnett's test for multiple comparisons.

### ClnR is a global regulator of gene expression in *C. difficile*

As antimicrobial resistance did not explain the changes in growth for the *clnR* and *clnAB* null mutants in LL-37, we hypothesized that there were changes in the expression of genes other than *clnRAB* in the *cln* mutants. To test this, we examined gene expression by RNA-seq for the *clnR* mutant grown with and without LL-37, compared to that of the parent strain (**Table S4**). This analysis revealed that 178 genes were differentially expressed at least 2-fold in the *clnR* mutant (*p* < 0.05). Notably, the *clnR* mutant demonstrated negative and positive changes in transcription, with many genes exhibiting additional conditional regulation by LL-37. In the absence of LL-37, the *clnR* mutant exhibited increased expression of 14 genes and decreased expression of 32 genes. Disruption of *clnR* had an even greater impact on expression in the presence of LL-37, resulting in increased expression of 29 genes and decreased expression of 103 genes. Of the 228 genes differentially regulated by LL-37 (**Table S1**), 56 were also influenced by *clnR* (**Table S5**). These results indicate that ClnR acts as both a repressor and inducer of gene expression, and that this regulatory potential is largely dependent on LL-37. The 178 genes regulated by ClnR fell into many different functional classes, with the most common ClnR-dependent genes encoding proteins with predicted metabolic functions (**Fig. 3**). These results support the premise that ClnR acts as a global transcriptional regulator in response to LL-37.

**Figure 3.**
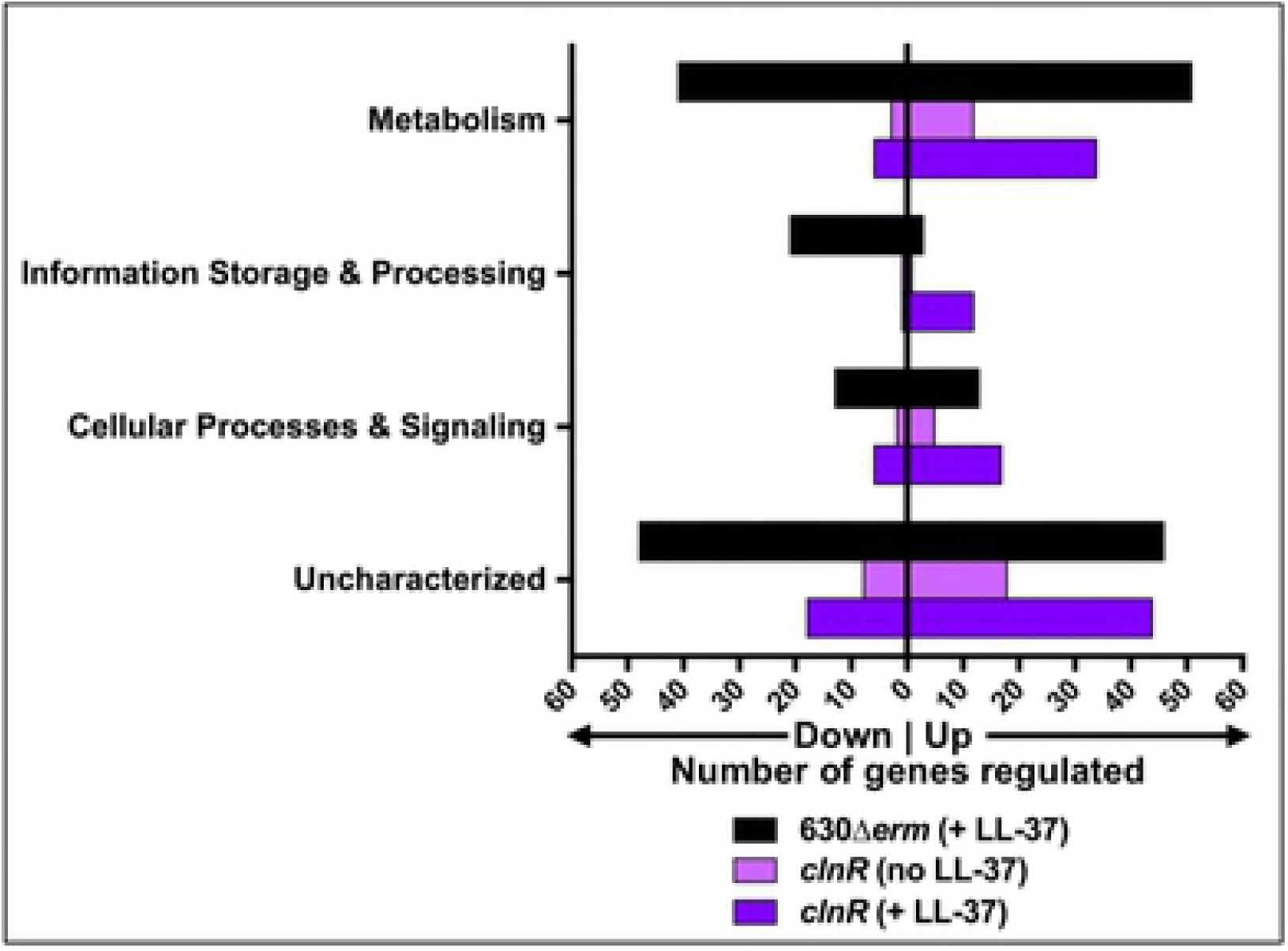
LL-37 and ClnR impact global gene expression. The genes listed in Table S1 (black) and Table S4 (light and dark purple) were assigned COG classifications according to the 2014 COG database. COG classifications were then grouped according to broader functions of Metabolism (COG classes C, E, F, G, H, I, P, and Q), Information Storage & Processing (COG classes A, B, J, K, and L), Cellular Processes & Signaling (COG classes D, M, N, O, T, U, V, W, Y, and Z), and Uncharacterized (COG classes R and S, or unassigned). Genes with functions that fall within two different groups are represented within both of the groups.

RNA-seq results from the *clnR* mutant and parent strain were validated by qRT-PCR for several apparent ClnR-dependent genes, as well as analysis of expression in the *clnAB* mutant and complemented strains (**Table S6**). Comparisons of *clnR* and *clnAB* mutant expression revealed that the regulator and transporter disruptions resulted in disparate effects on the transcription of some genes (*vanZ1, cdd4* and *iorA*) (**Table S6**). These disparate effects are most prominent in the presence of LL-37, highlighting that ClnR activity is dependent on LL-37. In some cases, the *clnR* and *clnAB* mutants had similar levels of expression, suggesting that the ClnAB transporter function is important for the activation of ClnR, whereas in other cases (*vanZ1, cdd4* and *iorA*) expression diverged in the *clnR* and *clnA* mutants, suggesting a role for the ClnAB transporter in the regulation of some LL-37-dependent genes, independent of ClnR. Complementation of *clnR* and *clnAB* was performed by restoring the entire *clnRAB* operon *in trans*, as restoration of expression was not possible using only the disrupted *clnR* or *clnAB*, respectively (**Fig. S2B**). As a result, the *clnR* complemented strain expresses the acquired copy of *clnRAB* and the native *clnAB.* Similarly, the *clnAB* complement expresses the native *clnR* and the acquired *clnRAB*. The altered ratios of components in these complement strains (**Fig. S2B**) may account for some discrepancies noted in these strains, further highlighting the complicated relationship between ClnR and ClnAB in regulation. These complex gene regulatory patterns suggest that multiple factors are involved in the transcription of some ClnR-regulated genes, and that ClnR has both direct and indirect effects on the expression of some loci.

### ClnR conditionally represses and induces *clnRAB* in response to LL-37

To understand the molecular mechanism of ClnR function, we further explored regulation of the *clnRAB* locus. ClnR is annotated as a GntR-family transcriptional regulator, and protein sequence comparisons suggest that it is a member of the YtrA sub-family of GntR regulators (**Fig. S4**). GntR-family regulators are most common among bacteria that inhabit complex environmental niches (29). The YtrA sub-family regulators are often found in conjunction with ABC-transporters and are typically autoregulatory (30). We examined the impact of *clnR* and *clnAB* disruption on expression of the *clnRAB* operon to determine if ClnR regulates expression of itself and *clnAB*. qRT-PCR analysis using primers located downstream of the *clnR* and *clnA* insertional disruptions revealed that transcription of *clnR* and *clnA* are increased in the *clnR* mutant grown without LL-37 (**Fig. 4**), suggesting that ClnR auto-represses the *cln* operon. Note that in the *clnR* insertional mutant, the *clnR* transcript cannot be translated into functional protein because of the insertional disruption, but *clnA* and *clnB* transcripts are produced and are expected to be translated. These results indicate that the disruption of *clnR* is not polar on expression of *clnA* (or *clnB*), but *clnAB* expression is disregulated in the *clnR* strain. In the presence of LL-37, *clnR* and *clnA* expression are no longer induced in the *clnR* mutant, demonstrating that ClnR activates the c*ln* operon in response to LL-37. Conversely, the *clnAB* mutant displays lower *clnR* and *clnA* expression during growth with or without LL-37 (**Fig. 4**). These results provide further evidence that the ClnAB transporter contributes to regulation of the *cln* operon and the ability of ClnR to respond to LL-37.

**Figure 4.**
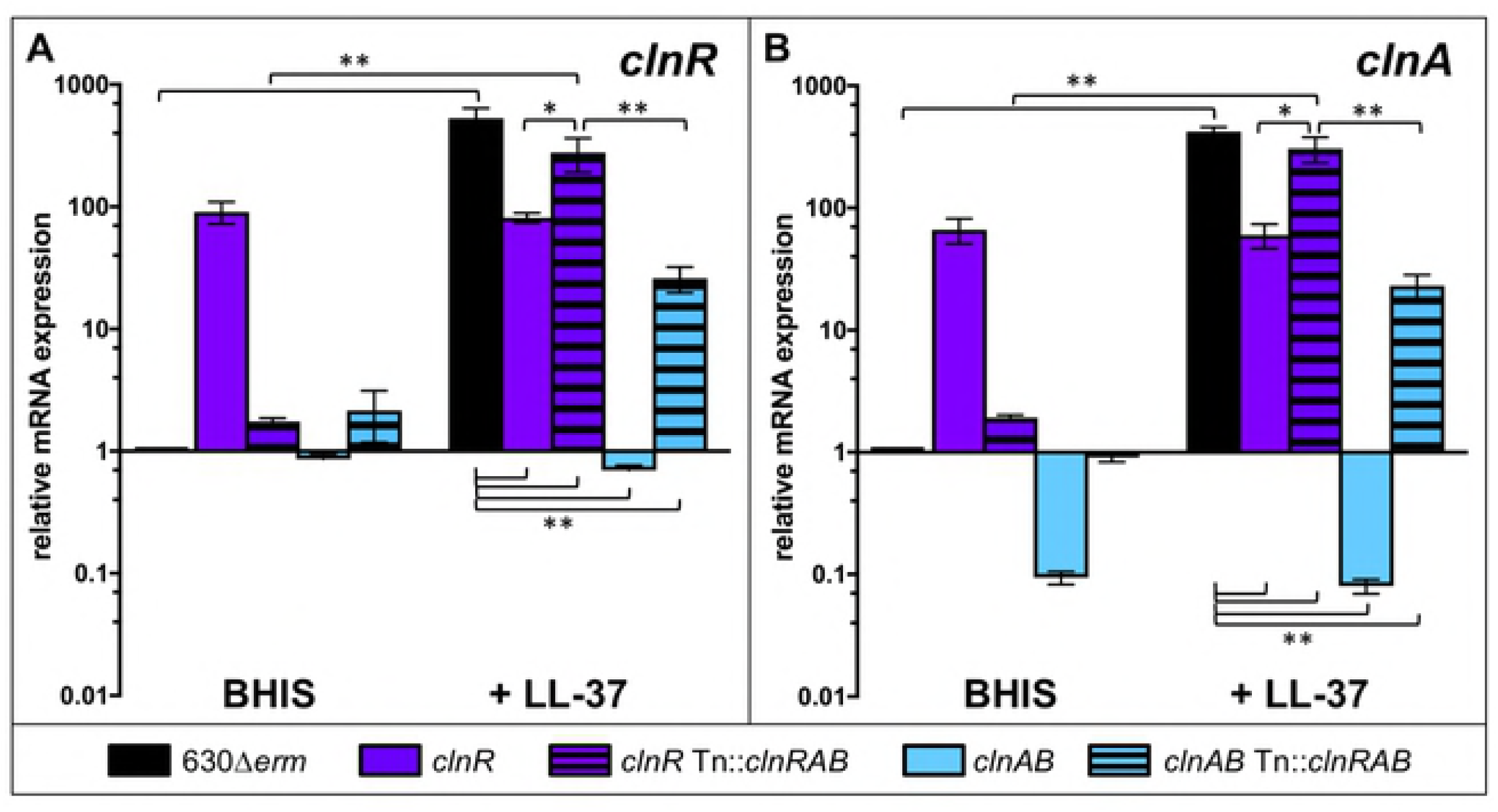
ClnR acts as a conditional repressor and inducer of *clnRAB* expression. Cultures of 630Δ*erm, clnR* (MC885)*, clnR* Tn::*clnRAB* (MC950)*, clnAB* (MC935), and *clnAB* Tn::*clnRAB* (MC953) were grown in BHIS alone or BHIS with 2 µg/ml LL-37. RNA samples were collected and processed for qRT-PCR analysis as described in Methods. Graphs show the mean mRNA expression levels of A) *clnR* and B) *clnA* relative to expression in strain 630Δ*erm* in BHIS alone. Error bars represent the standard error of the mean from at least three independent experiments. Data were analyzed by two-way ANOVA and Tukey’s multiple comparisons test, with comparisons indicated by brackets. * indicates *p* < 0.05, ** indicates *p* ≤ 0.0001.

### ClnR regulates the metabolism of different nutrient sources in *C. difficile*

Based on the evident changes in metabolic gene expression in the *clnR* mutant and the impact of LL-37 on ClnR-dependent transcription, we investigated the effect of ClnR and LL-37 on the growth of *C. difficile* with relevant metabolites. To this end, we grew the *clnR* mutant and the parent strain in minimal medium supplemented with glucose, fructose, mannose, mannitol, N-acetylglucosamine, or ethanolamine, with and without LL-37 (**Fig. 5**). We observed that for the first 2-3 h, growth of the *cln* mutants and the parent strain were indistinguishable in the presence or absence of LL-37, regardless of the supplemented carbon source (**Fig. 5**). Other groups have also observed preferential utilization of peptides by *C. difficile*, as amino acids are a preferred energy source for this bacterium (31-33). As anticipated, the addition of each of the examined carbon sources to minimal medium (MM) resulted in shorter doubling times (*i.e.,* faster growth) for the parent strain cultures (630Δ*erm*) than in the base minimal medium, with the exception of ethanolamine supplementation (**Fig. 5**, **Table S7,** column one). The *clnR* mutant grew less well than other strains in MM, MM with glucose, and MM with NAG, suggesting that ClnR is important for the utilization of peptides, glucose and NAG (**Fig. 5**). When a low concentration of LL-37 (0.5 µg/ml; 1/30 MIC) was added to the growth medium, the *clnR* and *clnAB* mutants grew better than the parent strain in base MM, MM with mannitol, MM with NAG, and MM with EA supplementation (**Fig. 5**), suggesting that ClnRAB is important for growth in a variety of nutrients in the presence of LL-37. These results indicate that the changes in growth observed with low levels of LL-37 are due to changes in ClnR-dependent bacterial metabolism, rather than the antimicrobial activity of LL-37. Moreover, the data strongly suggest that the growth and metabolism delays observed in LL-37 are mediated by ClnR through repression and activation of metabolic gene expression (**Tables S4, S5**).

**Figure 5.**
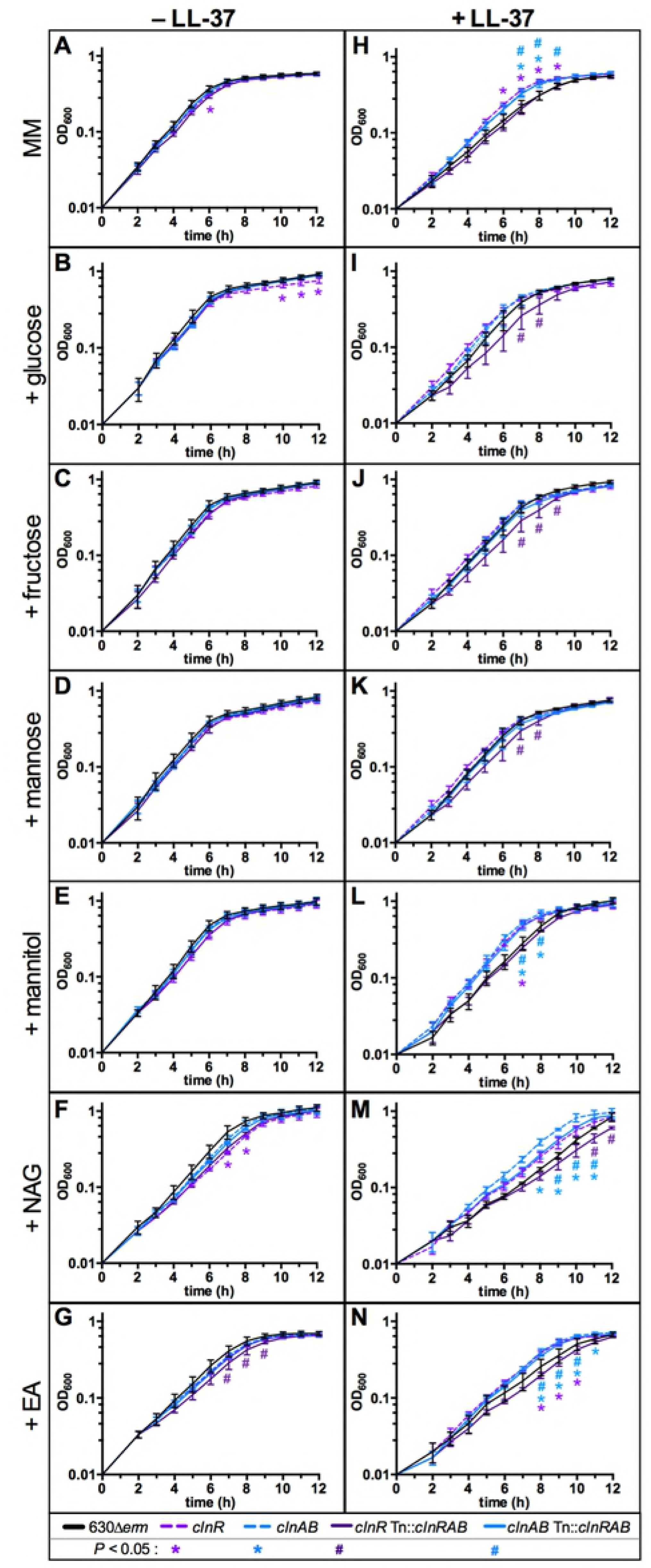
ClnR and LL-37 regulate the metabolism of nutrients. Active cultures of strain 630Δ*erm* (black), the *clnR* mutant (MC885, light purple), the *clnAB* mutant (MC935, light blue), *clnR* Tn::*clnRAB* (MC950, dark purple), and *clnAB* Tn::*clnRAB* (MC953, dark blue) were diluted to an OD_600_ in 0.01 in minimal media (MM) either without (A-G) or with (H-N) 0.5 µg/ml LL-37. Base minimal media (A, H) or MM supplemented with 30 mM glucose (B, I), 30 mM fructose (C, J), 30 mM mannose (D, K), 30 mM mannitol (E, L), 30 mM N-acetylglucoseamine (NAG) (F, M), or 30 mM ethanolamine (EA) (G, N). Graphs are plotted as the means +/− SEM from three independent replicates. Data were analyzed by two-way ANOVA with Dunnett’s multiple comparison test, comparing to 630Δ*erm* at each time point. Adjusted *p* value < 0.05 indicated by the symbols in the legend.

### ClnRAB modulates growth and virulence *in vivo*

As LL-37 is a host-produced peptide and *C. difficile* inhabits the gastrointestinal tract, the natural consequences of ClnR-LL-37–dependent gene regulation would appear during the growth of the pathogen in the host intestine. To examine the effects of *cln* mutants *in vivo*, we used the hamster and mouse models of CDI. Like humans, hamsters and mice are sensitive to colonization by *C. difficile*, especially after receiving antibiotics, and both produce cathelicidins similar to LL-37 (**Fig. S3**) (34-36). Syrian golden hamsters are acutely susceptible to infection by *C. difficile*, with as few as 1 to 10 CFU needed to produce fulminant disease (37). Mice are naturally not as susceptible to CDI as hamsters and require 10^4^-10^7^ CFU to achieve colonization, usually with low morbidity (38). For these reasons, the hamster model is most useful for examining early colonization and virulence, while mice allow for assessment of long-term *C. difficile* colonization (39).

Hamsters were infected with spores of 630*Δerm*, *clnR,* or *clnAB* strains and monitored for symptoms of infection as described in the Methods. Hamsters infected with either the *clnR* or *clnAB* mutant strains succumbed to infection more rapidly than animals inoculated with the parent strain, indicating that the *clnR* and *clnAB* mutants are more virulent (mean time to morbidity: 46.0 ± 12.2 h for 630Δ*erm*, 32.5 ± 5.8 h for *clnR*, 35.2 ± 6.1 h for *clnAB*; **Fig. 6A**). *C. difficile* disease is mediated by the two primary toxins, TcdA and TcdB. To determine whether the increased virulence of the *cln* strains was related to increased toxin levels, we extracted RNA from cecal samples collected from animals at the time of morbidity and performed digital droplet PCR for absolute quantification of *tcdA*, *tcdB*, and *clnR* expression (**Fig. S5**). No significant differences in toxin expression were apparent at the time of morbidity. Because only one timepoint could be assessed, the results do not resolve whether the *clnR* and *clnA* mutants had altered toxin expression during the course of infection. But, the time from infection to morbidity for *clnR* and *clnAB* infections indicate that these mutants produce toxin earlier in the course of infection, resulting in earlier symptoms of disease and morbidity.

**Figure 6.**
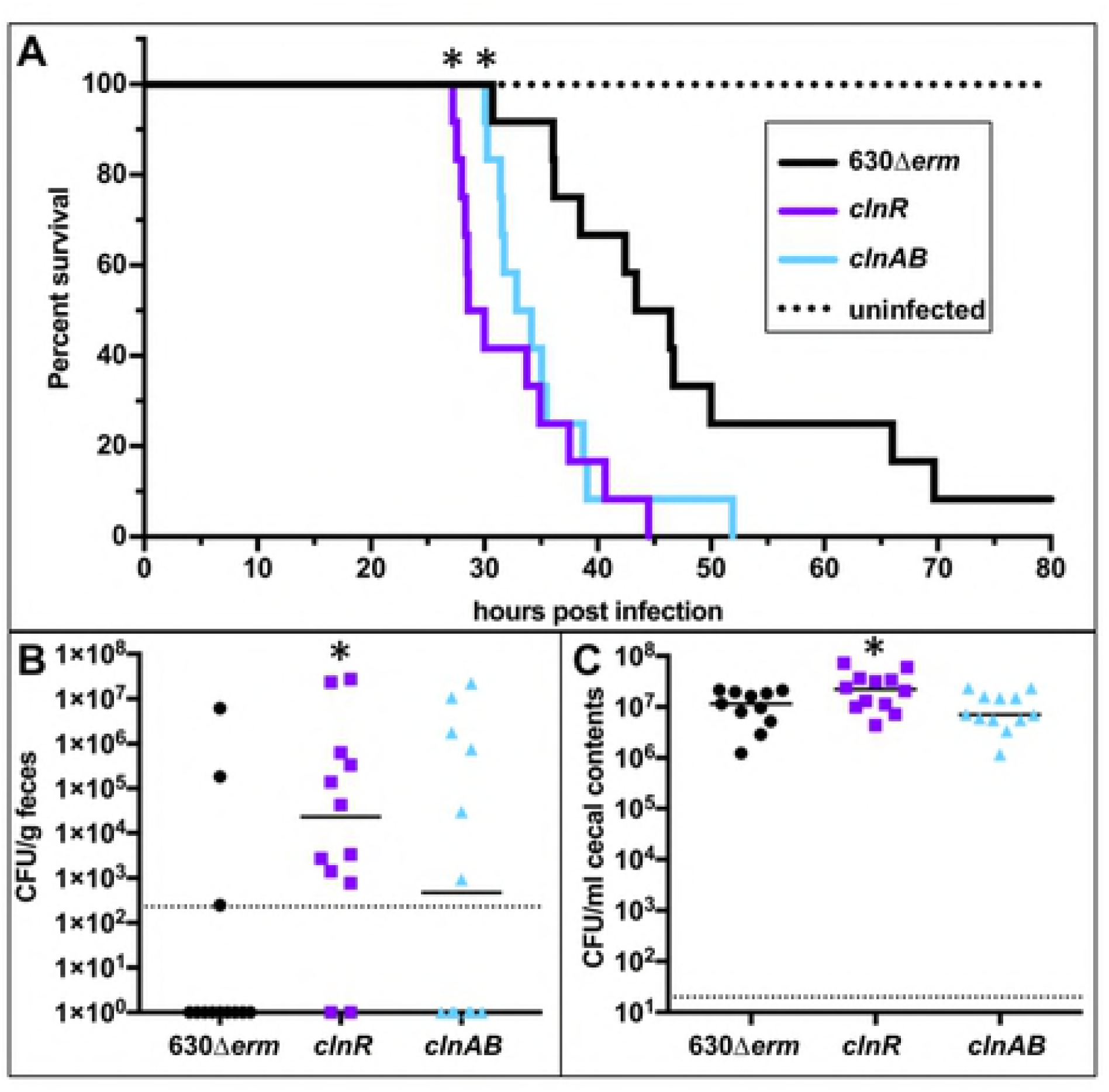
*clnR* and *clnAB* mutants are more virulent in a hamster model of infection. Syrian golden hamsters were inoculated with approximately 5000 spores of strain 630Δ*erm* (n = 12), *clnR* (MC885; n = 12), or *clnAB* (MC935; n = 12). A) Kaplan-Meier survival curve depicting time to morbidity. Mean times to morbidity were: 630Δ*erm* 46.0 ± 12.2 (n=11); *clnR* 32.5 ± 5.8 (n=12); *clnA* 35.2 ± 6.1 (n=12). * indicates *p* ≤ 0.01 by log-rank test. B) Total *C. difficile* CFU recovered from fecal samples collected at 12 h.p.i. Dotted line demarcates lowest limit of detection. Solid black line marks the median. Fisher’s exact test compared the number of animals without and with detectable CFU compared to 630Δ*erm* (* indicates *p* < 0.05). C) Total *C. difficile* CFU recovered from cecal contents collected post-mortem. Dotted line demarcates limit of detection. Solid black line marks the median. Numbers of CFU are compared to 630Δ*erm* by one-way ANOVA with Dunnett’s multiple comparisons test (* indicates *p* < 0.05).

To assess *C. difficile* colonization by the different strains, hamster fecal samples were taken at 12 h post-infection and plated onto selective medium. *C. difficile* was recovered from fecal samples in significantly more animals infected with the *clnR* strain than in the 630Δ*erm*-infected group, suggesting that the *clnR* mutant colonizes the hamster intestine more rapidly (**Fig. 6B**). In addition, the *clnR* mutant reached a higher bacterial burden at the time of morbidity (1.2 × 10^7^ CFU/ml for 630Δ*erm*, 2.7 × 10^7^ CFU/ml for *clnR*; **Fig. 6C**). These results illustrate that the *clnRAB* operon plays a significant role in the colonization and virulence dynamics of *C. difficile* hamster infections.

The colonization results in hamsters suggested a role for ClnRAB in colonization dynamics, which was further examined in the mouse model. Mice were infected with spores of either 630Δ*erm*, *clnR*, *clnAB*, *clnR* Tn::*clnRAB*, or *clnAB* Tn::*clnRAB* strains and monitored for colonization and disease as described in the Methods. Mice infected with *clnR* lost less weight and recovered more quickly than mice infected with 630Δ*erm* (**Fig. 7A**). Additionally, mice infected with *clnR* cleared the bacteria more quickly, with fewer animals having detectable CFU in their feces (**Fig. 7B; Fig. S6**). While the impact on short-term colonization and virulence in hamsters and longer-term colonization and persistence in mice are contrasting, the results from both animal models strongly suggest that ClnR contributes to the ability of *C. difficile* to initiate colonization, cause disease, and persist in the intestinal environment.

**Figure 7.**
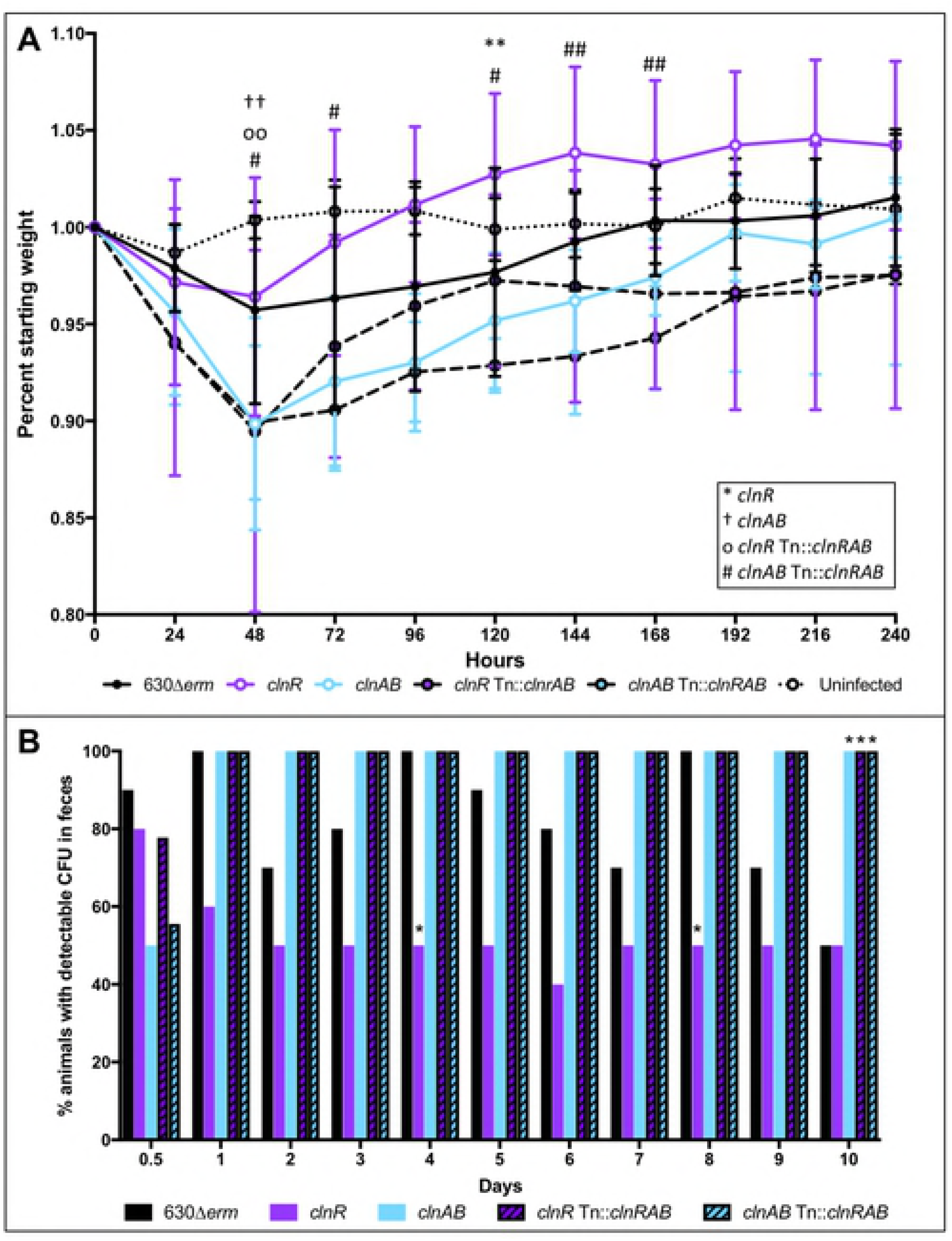
Mice infected with a *clnR* mutant recover more quickly. C57BL/6 mice were inoculated with approximately 1 × 10^5^ spores of 630Δ*erm* (n = 10), *clnR* (MC885; n = 10), *clnAB* (MC935; n = 10), *clnR* Tn::*clnRAB* (MC950; n = 9), or *clnAB* Tn::*clnRAB* (MC953; n = 9) as detailed in Methods. A) Daily weights were calculated as a percent of the animal’s starting weight. Graph shows the mean ± SEM. Data were analyzed by two-way ANOVA with Dunnett’s multiple comparisons test, comparing each strain to 630Δ*erm*. See legend for significance symbol key. One symbol indicates adjusted *p* value <0.05, two symbols indicates *p* <0.002. B) The percentage of animals with detectable CFU in their feces (Days 0.5 **–** 9) or cecal contents (Day 10). Data were analyzed by one-way ANOVA with Dunnett’s multiple comparisons test comparing each strain to 630Δ*erm* at the same time point. * indicates adjusted *p* value <0.05.

Considering that differences in either sporulation or germination rates can also influence virulence and bacterial burden i*n vivo,* we assessed sporulation and germination for the *clnR* and *clnAB* mutants for defects in either process. No significant difference in sporulation or germination rates was observed for either mutant (**Fig. S7**).

### ClnRAB and LL-37 promote toxin production

Since toxin production is the primary virulence factor leading to *C. difficile* symptoms, we further investigated the effects of LL-37 and ClnRAB on toxin production under more controlled conditions *in vitro*. qRT-PCR analysis of *tcdA* and *tcdB* transcription was assessed for the *cln* mutants and parent strain during logarithmic phase growth in BHIS medium, with or without added LL-37. As shown in **Fig. S8A, B**, LL-37 exposure resulted in increased expression of *tcdA* (4.6-fold) and *tcdB* (2.2-fold) in wild-type cells. In contrast, the *clnR* and *clnAB* mutants demonstrated lower expression of toxins in LL-37, suggesting that ClnRAB is partially responsible for LL-37-dependent regulation of toxin expression *in vitro*. Toxin expression is known to be controlled by several regulatory factors, many of which respond to low nutrient availability and/or the transition to stationary phase growth (40). To determine which of the toxin regulators may be influenced by LL-37, we examined expression of regulators and regulator-dependent factors, including *tcdR*, *sigD*, *ilcV* (as an indicator of CodY activity), and *CD0341* (as an indicator of CcpA activity) (**Table S8**). Of these, only *ilvC* expression is statistically altered in LL-37; however, the increase in *ilvC* expression is far more modest (2.2-fold increase) than would be expected with robust CodY activation (18, 19).

Because toxin production is typically low at mid-logarithmic phase in BHIS medium, we also examined toxin protein levels after 24 h growth in TY medium. Western blot analysis indicated that TcdA levels were significantly lower in the *clnR* mutant in TY medium, relative to the parent strain. When cells were grown in medium supplemented with LL-37, final TcdA levels decreased about 3-fold in the parent strain (**Fig. S8C**). In contrast, TcdA levels did not change for the *clnR* and *clnAB* mutants in LL-37. While these findings contradict the induction of *tcdA* expression observed at log-phase in BHIS medium, the data support the observation that LL-37 and ClnRAB influence toxin expression, and that the outcome of this regulation on toxin production is dependent on growth conditions. These observations provide further evidence that the ClnRAB system is involved in toxin production and that this system is necessary for the influence of LL-37 on toxin production.

### ClnR acts as a DNA-binding regulator that binds multiple promoters

As a predicted GntR-family transcriptional regulator, we hypothesized that ClnR binds DNA. Because expression results suggested that ClnR is autoregulatory (**Fig. 4**), we initially tested whether ClnR directly regulates its own promoter. We produced recombinant His-tagged ClnR and performed gel shifts with fluorescein-labeled DNA of the 84 bp upstream of the predicted *clnR* transcriptional start site. This DNA fragment was selected because it encompasses a predicted σ^A^-dependent promoter with −10 (at −52 to −47 bp) and −35 (at −73 to −68 bp) consensus sequences and a tandem repeat sequence (at −46 to −16 bp) that includes a possible ClnR-binding site (**Fig. S9**). Incubation of His-ClnR with this DNA fragment resulted in a shift visible after electrophoresis, both with and without LL-37 (**Fig. 8A**). The apparent *K*_d_ value for this interaction was calculated to be 118 nM (± 40 nM) without LL-37 and 85 nM (± 7 nM) with LL-37, indicating that the affinity of ClnR for this DNA sequence does not change significantly in the presence of LL-37 in these conditions.

**Figure 8.**
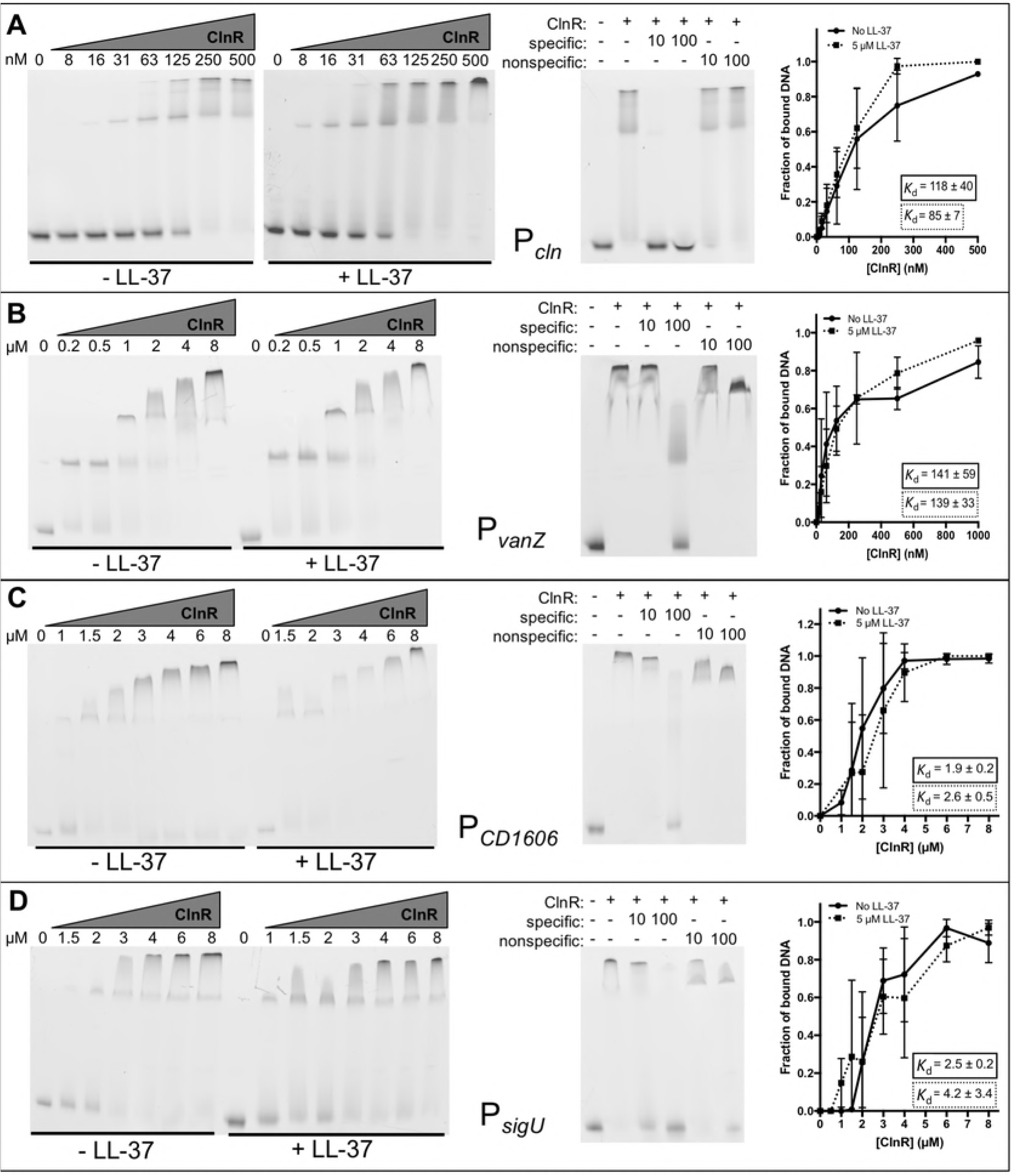
ClnR directly and specifically binds several DNA targets. Electrophoretic mobility shift assays were performed as described in Methods using His-tagged ClnR and fluorescein-labeled DNA encompassing regions upstream of A) *clnR,* B) *vanZ,* C) *CD1606,* or D) *sigU*. ClnR was added to reactions at varying concentrations (specified in nM in A, in µM elsewhere) either without or with 0.5 µM LL-37, as indicated. Competitive EMSAs were performed with the addition of unlabeled target DNA (specific) or unlabeled P*spo0A* DNA (non-specific) at either 10x or 100x the concentration of labeled target DNA. 125 nM ClnR was used for the competitive EMSA for P*cln,* 8 µM for all others. Apparent *K*_d_ values were calculated as described in Methods. Graphs are the binding curves showing the mean and standard deviation from three independent replicates.

Additional ClnR-regulated promoters were examined for direct binding, including predicted upstream promoter elements for the metabolic operons *grd* (CD630_23540), *mtl* (CD630_23340), and *ior* (CD630_23810); other transcriptional regulators, including *sigU* (*csfU, CD630_18870*) and *CD630_16060*, as well as the uncharacterized *vanZ* ortholog (CD630_12400). ClnR bound to all of these promoter sequences but exhibited specificity for P*vanZ*, P*CD1606*, and P*sigU*, with or without LL-37 (**Fig. 8B-D**). Binding was less specific for P*grd,* P*mtl* and P*ior* under the conditions tested (**Fig. S10**). The calculated apparent *K*_d_ for P*vanZ* was 141 nM (± 59 nM) without LL-37 and 139 nM (± 33 nM) with LL-37, the apparent *K*_d_ for P*CD630_16060* was 1.9 µM (± 0.2 µM) without LL-37 and 2.6 µM (± 0.5 µM) with LL-37, and the apparent *K*_d_ for P*sigU* was 2.5 µM (± 0.2 µM) without LL-37 and 4.2 µM (± 3.4 µM) with LL-37 (**Fig. 8**). Because the apparent *K*_d_ values for these targets are similar both with and without LL-37, it does not appear that LL-37 influences ClnR binding of these targets in these *in vitro* conditions.

### ClnR directly binds LL-37

Although the EMSA did not uncover differences in ClnR-DNA binding in the absence or presence of LL-37 *in vitro* for the DNA targets examined, our expression and growth data suggest that ClnR regulates transcription in response to LL-37. Based on *clnR* mutant phenotypes for cells grown with and without LL-37, we hypothesized that ClnR directly binds LL-37 to regulate ClnR activity. To test this hypothesis, we performed surface plasmon resonance (SPR) to examine the binding kinetics of ClnR with LL-37. These experiments showed that His-ClnR interacts with LL-37 with a *K*_d_ of 83 ± 14 nM (**Fig. 9A**). This *K*_d_ value is similar to the apparent *K*_d_ values that were calculated for the affinity of His-ClnR for P*cln* DNA. This result suggests that the concentrations of ClnR needed to interact with both LL-37 and P*cln* DNA are similar, and that interactions of these three components could occur simultaneously. Furthermore, the interaction between His-ClnR and LL-37 is specific, as SPR using scrambled LL-37 found no apparent interaction between these molecules (**Fig. 9B**).

**Figure 9.**
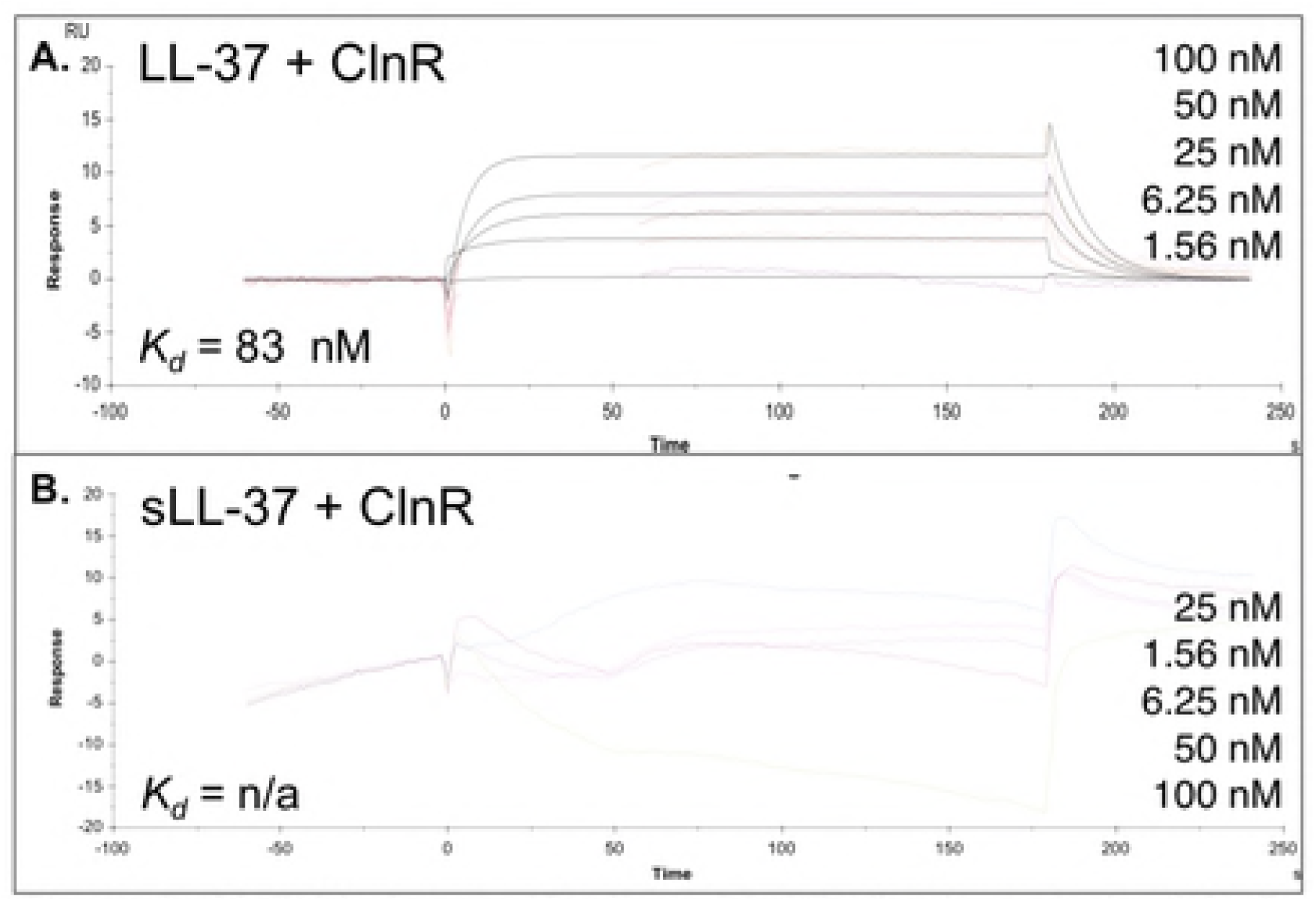
ClnR binds specifically to LL-37. Kinetic analysis of ClnR-LL-37 binding. Surface Plasmon Resonance (SPR) sensorgram quantifying binding of His-ClnR to immobilized **A)** LL-37 or **B)** scrambled LL-37 (sLL-37; negative control) measured as the relative response in resonance units (RU) after reference background subtraction. The concentrations of His-ClnR applied are listed in their order on the graphs, respectively. Affinity modeling (1:1) was used to calculate the dissociation constant for binding. LL-37-ClnR: *Kd* = 83 ± 14 nM; sLL-37-ClnR: no significant binding detected

## DISCUSSION

Many bacteria encode signaling systems for detecting conditions within the host environment, allowing for activation of genes that are necessary for survival within the host. LL-37 acts as a signal for many pathogens to adapt to the host, though most of the mechanisms that have been investigated are implicated in bacterial virulence and antimicrobial resistance (41-47). Little is known about the molecular mechanisms that *C. difficile* uses to adapt and survive in the host intestinal environment. Our results have revealed that LL-37 alters global gene expression in *C. difficile* through the previously unknown regulator and ABC-transporter system, ClnRAB. Moreover, the activation of ClnRAB is specific to LL-37 and independent of the antimicrobial effects of this host peptide.

Many of the genes regulated by LL-37 and ClnRAB function in metabolism and energy production (**Table S4, Fig. 3**). The regulation of metabolic pathways in response to LL-37 was previously observed in other pathogens, including *S. pyogenes*, *E. coli, P. aeruginosa*, and *S. pneumonia*e, but their role in the bacterial response to LL-37 has not been clear (43, 46-48). Our data indicate that the regulation of genes by LL-37/ClnRAB *in vivo* has notable effects on *C. difficile* colonization and virulence (**Fig. 6, Fig. 7**). Results from the mouse infection model suggest that disruption of ClnRAB results in dysregulation of metabolism that significantly hinders the ability of *C. difficile* to colonize; whereas in the exquisitely toxin-sensitive hamster model, the metabolism defects in *cln* mutants quickly progress to nutrient deprivation and toxin production. These effects are not unexpected, given that nutrient deprivation is demonstrably the primary factor driving *C. difficile* toxin expression (20, 32, 40, 49-54). *C. difficile* possesses an unusual metabolic repertoire for energy generation, including solventogenic fermentation (55, 56), Stickland (amino acid) fermentation (31), and autotrophic growth via the Wood-Ljungdahl pathway (57). However, the importance of most of these individual metabolic pathways for growth and virulence *in vivo* has not been determined. Because ClnR is a global regulator that negatively and positively influences the expression of multiple metabolic pathways, many of which are constitutively expressed in a *clnR* mutant, we cannot infer which of these pathways are most influential for host pathogenesis. Determining how and which ClnR-controlled pathways and mechanisms influence disease could expose potential vulnerabilities of *C. difficile* that may be exploited to prevent infections.

Overall our results indicate that ClnRAB responds directly and specifically to LL-37 without conferring LL-37 resistance and suggest that ClnR responds to LL-37 as an indicator of the host environment, conferring a colonization advantage. The *clnRAB* locus is highly conserved in *C. difficile,* with representation at ≥ 99% amino acid sequence identity in over 500 strains at the time of this publication (NCBI, BLASTp). The data strongly suggest that ClnR acts as a pleiotropic regulator in *C. difficile* that controls the expression of genes involved in metabolism and virulence, in response to the host peptide, LL-37. Further study of the activation and downstream impacts of this regulatory pathway will contribute greatly to our understanding of how *C. difficile* adapts to the host environment and causes disease.

## METHODS

### Bacterial strains and growth conditions

**Table S8** lists the bacterial strains and plasmids used in this study. *Escherichia coli* was grown aerobically in LB medium (Teknova) at 37°C (58). As needed, cultures were supplemented with 20 µg chloramphenicol ml^−1^ (Sigma-Aldrich) or 100 µg ampicillin ml^−1^ (Cayman Chemical Company). *C. difficile* was grown at 37°C in an anaerobic chamber containing 10% H_2_, 5% CO_2_ and 85% N_2_ (Coy Laboratory Products) in brain heart infusion medium supplemented with 2% yeast extract (BHIS; Becton, Dickinson, and Company), TY broth (51), or 70:30 sporulation agar (59), as previously described (60). As needed, *C. difficile* cultures were supplemented with LL-37 (Anaspec) at the concentrations stated in the text, or 2 µg thiamphenicol ml^−1^ (Sigma-Aldrich) for plasmid selection.

### Strain and plasmid construction

Plasmids and strains used in this study are listed in **Table S9**. **Table S10** lists the oligonucleotides used in this study. *C. difficile* strain 630 (GenBank accession NC_009089.1) served as the reference for primer design and cloning. PCR amplification was performed using genomic DNA from strain 630Δ*erm* as a template. PCR, cloning, and plasmid DNA isolation were performed according to standard protocols (60-62). Plasmids were confirmed by sequencing (Eurofins MWG Operon).

Null mutations in *C. difficile* genes were introduced by re-targeting the group II intron of pCE240 using the primers listed in **Table S10,** as previously described (16, 61). Plasmids were introduced into *E. coli* strain HB101 pRK24 via transformation. *E. coli* strains were then conjugated with *C. difficile* for plasmid transfer. Transconjugants were exposed to 50 µg kanamycin ml^−1^ to select against *E. coli*, 10 µg thiamphenicol ml^−1^ to select for plasmids, and subsequently, 5 µg erythromycin ml^−1^ to select for insertion of the group II intron into the chromosome. Insertion of the group II intron into erythromycin resistant clones was confirmed by PCR using the primers listed in **Table S10**.

The *clnRAB* coding sequence and apparent promoter (pMC649) were introduced into the *Tn*916 transposon of BS49 as MC951 (20). MC951 was then mated with *C. difficile* strains MC885 and MC935 to generate MC950 and MC953, respectively. Transconjugants were exposed to 50 µg kanamycin ml^−1^ to select against *B. subtilis* and 5 µg erythromycin ml^−1^ to select for integration of the transposon. Insertion of the genes was confirmed by PCR using the primers listed in **Table S10**. Complete information on plasmid construction is available in the supplemental materials (**Table S11**).

### RNA sequencing (RNA-seq)

Active cultures of 630Δ*erm* or the *clnR* mutant were diluted to approximately OD_600_ 0.05 in BHIS alone or with 2 µg LL-37 ml^−1^ and grown to OD_600_ 0.5 for harvesting. RNA was extracted and DNase I treated as previously described (18, 63). rRNA was depleted from the total RNA using the Bacterial Ribo-Zero™ rRNA Removal Kit (EpiCentre, Madison, USA) following the manufacturer’s instructions. cDNA libraries were prepared with the ScriptSeq v2 RNA-Seq library preparation kit (Epicentre, Madison, USA). Briefly, the rRNA depleted sample was fragmented using an RNA fragmentation solution prior to cDNA synthesis. The fragmented RNA was further reverse transcribed using random hexamer primers containing a tagging sequence at their 5′ ends, 3′ tagging was accomplished using the Terminal-Tagging Oligo (TTO). The di-tagged cDNA was purified using the AMPure™ XP (Agencourt, Beckmann-Coulter, USA). The di-tagged cDNA was further PCR amplified to add index and sequencing adapters, the amplified final library was purified using AmpureXP beads. The final pooled libraries were sequenced on the Illumina HiSeq3000 system in a Single-end (SE) 150 cycle format, each sample was sequenced to approximate depth of 8-12 million reads. Sequenced reads were aligned to the CD630Derm (GenBank Accession GCA_000953275.1) genome reference for the 630Derm strain of *C. difficile* using the STAR Aligner (version 2.4.0g1; (64)). Counts of reads that uniquely map to genes in the reference genome annotation were accumulated using htseq-count (HTSeq 0.6.1p1; (65)). Samples from two independent experiments were library size normalized separately in DESeq2 (66) and the resulting normalized gene read counts were used as the gene abundance estimation and imported into Excel for gene expression comparisons. Gene abundances from the two experiments were averaged, and data were analyzed using the Student’s two-tailed t-test. Cluster of orthologous genes (COG) designations were assigned according to the NCBI COG database (2014 updated version) (67). Sample preparation and analyses were performed by the Yerkes Nonhuman Primate Genomics Core (Emory University). Raw data files are available in the NCBI-SRA database under accession number (pending).

### Quantitative reverse transcription PCR analysis (qRT-PCR)

Active cultures were diluted to an OD_600_ of approximately 0.05 in BHIS alone or BHIS with antimicrobials. The antimicrobials used included: LL-37 (Anaspec), scrambled LL-37 (Anaspec), mCRAMP (Anaspec), SMAP-29 (Anaspec), ampicillin (Cayman Chemicals), vancomycin (Sigma Aldrich), nisin (MP Biomedicals), or polymyxin B (Sigma Aldrich). Cultures were harvested at an OD_600_ of 0.5, mixed with 1:1 ethanol:acetone on ice and stored at −80°C. RNA was extracted, DNase I treated, and used to generate cDNA as described above for RNA sequencing. qRT-PCR reactions were performed using the Bioline Sensi-Fast SYBR and Fluroescein kit on a Roche LightCycler 96 instrument. Primers were designed with the assistance of the IDT PrimerQuest tool (Integrated DNA Technologies) and are listed in **Table S10**. Each qRT-PCR reaction was performed as technical triplicates for at least three biological replicates. The ΔΔC_t_ method was used to normalize expression to *rpoC,* an internal control transcript, for relative quantification (68). Statistical analysis of the results was performed using GraphPad Prism version 7 for Macintosh (GraphPad Software, La Jolla, GA) to perform either one- or two-way analysis of variance (ANOVA) with Dunnett’s or Sidak’s multiple-comparison test, as indicated.

### Droplet Digital PCR (ddPCR)

RNA was extracted from cecal samples as previously described (22). RNA was subsequently DNase I treated and used to generate cDNA as described above for qRT-PCR. cDNA was diluted to a final concentration of 5 ng/µl RNA equivalent. Samples were prepared in duplicate with 1.25 ng/µl cDNA, 70 nM each of forward and reverse primers (as listed in **Table S10**), and 1x QX200 ddPCR EvaGreen Supermix (Bio-Rad). 20 µl of each sample was loaded into a Bio-rad DG8 cartridge for droplet generation in a Bio-Rad QX200 Droplet Generator with 70 µl Droplet Generation Oil for EvaGreen (Bio-Rad) per sample. Droplets were transferred to an Eppendorf Twin-Tech 96-well plate, which was sealed with foil prior to PCR on a C1000 Touch thermal cycler with the following reaction parameters: 5 min at 95°C, 40 rounds of 30 s at 95°C and 1 min at 53°C, 5 min at 4°C, 5 min at 90°C (all steps with 2°C/s ramp). Droplets were then read on the Bio-Rad QX200 Droplet Reader. Samples without reverse transcriptase were run as a negative control and were used as reference to manually set the threshold values for positive calls in the QuantaSoft analysis software. Samples were only analyzed for *tcdA*, *tcdB,* and *clnR* expression if *rpoC* transcripts were detected (as a housekeeping gene, the detection of *rpoC* indicates sufficient *C. difficile* genomic material was present in the sample). Statistical analysis of the results using GraphPad Prism version 7 for Macintosh (GraphPad Software, La Jolla, GA) to perform two-way ANOVA with Dunnett’s multiple-comparison test.

### Minimum inhibitory concentration (MIC) and minimum bactericidal concentration (MBC)

Minimum inhibitory concentrations (MIC) were determined as previously described (69). MICs were determined for LL-37 (Anaspec), ampicillin (Cayman Chemicals), vancomycin (Sigma Aldrich), nisin (MP Biomedicals), and polymyxin B (Sigma Aldrich). Briefly, overnight cultures of *C. difficile* strains were diluted 1:50 in Mueller-Hinton Broth (Difco) and grown to OD_600_ of 0.45 (∼5 × 10^7^ CFU/ml). Cultures were then diluted 1:10 in MHB and seeded at a further 1:10 dilution in a round-bottom 96-well plate prepared with serial dilutions of antimicrobials for a starting concentration of ∼5 × 10^5^ CFU/ml. Plates were incubated for 24 h at 37°C in the anaerobic chamber. The MIC was determined as the lowest concentration of antimicrobial at which no growth was visible after 24 h. For MBC determination, the full volume of wells at concentrations at and above the MIC were transferred as a 1:10 dilution into BHIS and incubated for 24 h at 37°C in the anaerobic chamber. The minimum bactericidal concentration (MBC) was determined as the lowest concentration of antimicrobial at which no growth was visible after 24 h.

### Western blots

*C. difficile* strains were grown in BHIS medium containing 0.2% fructose and 0.1% taurocholate, as previously described (20). Cultures were diluted into BHIS medium and grown to OD_600_ of ∼0.5, then diluted 1:10 into TY medium with or without 2 µg LL-37 ml^−1^ and grown for 24 h at 37°C. Cells were harvested by centrifugation, resuspended in SDS-PAGE loading buffer (without dye) and mechanically disrupted as previously described (19, 70). Protein concentrations were assessed using a micro BCA assay (Thermo Scientific) and 6 µg of whole cell protein was loaded onto a 12% polyacrylamide gel (Bio-Rad). Proteins were subsequently transferred from the SDS-PAGE gel onto nitrocellulose membranes (0.45 µM; Bio-Rad), and probed with mouse anti-TcdA antibody (Novus Biologicals). Membranes were then washed and probed with goat anti-mouse secondary Alexa Fluor 488 antibody (Life Technologies). Imaging and densitometry analyses were performed using a ChemiDoc MP and Image Lab Software (Bio-Rad). Density of the TcdA band was normalized to total protein density. Three biological replicates were analyzed for each strain and condition. Statistical analyses were performed using either a Student’s *t* test with Holm-Sidak correction or a one-way ANOVA, followed by a Dunnett’s multiple comparisons test.

### Electrophoretic mobility shift assays (EMSAs)

Recombinant N-terminally His-tagged ClnR was produced by GenScript (Piscataway, NJ). Gene transcription in the *clnR* mutant complemented with His-tagged ClnR confirmed the functionality of this protein (**Table S12**). 5’-fluorescein-labeled DNA (10 ng per reaction; purified by extraction from a 4-20% TGX polyacrylamide gel) was incubated for 30 min at 37°C with His-ClnR (0 – 8 µM) with 10 mM Tris, pH 7.4, 10 mM MgCl_2_, 100 mM KCl, 7.5% glycerol, and 2mM DTT. 50 ng of salmon sperm DNA was added to each reaction as a noncompetitive inhibitor. In competition experiments, either 100 ng (10x) or 1 µg (100x) unlabeled target DNA (specific) or unlabeled P*spo0A* (nonspecific) DNA was incubated with His-ClnR (125 nM for P*cln* reactions, 8 µM for other targets) for 20 min at 37°C prior to the addition of labeled P*clnR* DNA for a further 10 min incubation. Reactions were loaded onto a pre-run 4-20% TGX polyacrylamide gel (Bio-Rad) and imaged on a Typhoon phosphoimager (GE Lifesciences) using the 520 BP fluorescence channel. Images from at least three replicates were analyzed in ImageLab (Bio-Rad) to determine the density of signal in bound and unbound fractions. Using GraphPad Prism, apparent K_d_ values were calculated by non-linear regression using an equation for cooperative binding of Y = F_max_*((x/*K*_d_)^n^)/(1+(x/*K*_d_)^n^), where Y = the fraction of bound DNA, x = the concentration of ClnR, F_max_ = the saturation level of bound DNA, *K*_d_ = the concentration of ClnR when half of the DNA is bound, and n = cooperativity coefficient (71).

### Growth curves in minimal medium

Growth curves were performed using a minimal medium based on a previously described complete defined minimal media (CDMM), but lacking D-glucose as used by Cartman *et al.* and adjusted to pH 7.4 (33, 72). The base medium was supplemented with 30 mM D-glucose (Sigma-Aldrich), 30 mM D-fructose (Fisher), 30 mM D-mannose (BD Difco), 30 mM D-mannitol (Amresco), 30 mM N-acetylglucosamine (Chem-Impex), or 30 mM ethanolamine-HCl (Sigma-Aldrich), as noted. Growth curves in minimal medium (MM) were carried out as follows: log-phase cultures were grown to an OD_600_ of 0.5 in BHIS medium, then diluted 5-fold into MM. Diluted cultures were then used to inoculate minimal medium broth for growth assays at a starting OD_600_ of ∼0.01 (2 ml into 23 ml of MM). Growth curve data were analyzed by two-way ANOVA with Dunnett’s test for multiple comparisons, comparing each strain to 630Δ*erm* at each time point. Doubling times were analyzed by one-way ANOVA with Dunnett’s test for multiple comparisons or by Student’s *t* test, as indicated.

### Surface Plasmon Resonance (SPR)

Using a Biacore X100 instrument (GE Healthcare), LL-37 (10 µm in acetate buffer, pH 5.0; Anaspec) was immobilized onto a CM5 sensor chip (GE Healthcare) using standard amine-coupling at 25°C and targeted at 2000 RU (actual RU 2080) (73). sLL37 (Anaspec) was immobilized on a separate CM5 sensor chip using the same procedure (final signal RU 1726). Running buffer I was HBS-P buffer (GE Healthcare). Running buffer II was composed of 113 mM NaCl, 24 mM NaHCO_3_, 3.9 mM KCl, 1.3 mM CaCl_2_, 0.6 mM MgCl_2_, 0.005% surfactant P20, pH 7.3 (74). His-ClnR (as described above for EMSAs) was dialyzed into running buffer II, then diluted into running buffer II for two-fold serial dilutions from 10 µM to 0.625 µM. His-ClnR was injected over immobilized LL-37 or sLL-37 for 180 s association time and 300 s dissociation time. The chip surfaces were regenerated by injecting 1 M NaCl in 50 mM NaOH for 90 s, then the surface was stabilized for 300 s prior to the next cycle run. The flow rate used was 10 µl/min. Experiments were performed a minimum of two times.

### Animal studies

Male and female Syrian golden hamsters (*Mesocricetus auratus;* Charles River Laboratories) were housed individually in sterile cages within a biosafety level 2 facility in the Emory University Division of Animal Resources. Sterile water and rodent feed pellets were available for the animals to consume *ad libitum*. Hamsters were administered 30 mg/kg body weight clindamycin (Hospira) by oral gavage 7 days prior to inoculation with *C. difficile*, to promote susceptibility to infection (19, 70). Spores were prepared as previously described (35, 36) and stored in phosphate-buffered saline (PBS) with 1% bovine serum albumin. Spores were heated for 20 minutes at 60°C and cooled to room temperature prior to inoculating hamsters. Hamsters were administered approximately 5,000 spores of strains 630Δ*erm*, *clnR* (MC885), or *clnAB* (MC935) by oral gavage and monitored for signs of disease. Hamsters were considered moribund after ≥ 15% weight loss from maximum body weight or when lethargic, with or without concurrent diarrhea and wet tail. Hamsters were euthanized once reaching either of these criteria. Fecal samples were collected daily, and cecal samples were collected post-mortem at the time of morbidity. Colony forming units (CFU) in fecal and cecal samples were plated on TCCFA medium as described previously (20). Differences in CFU counts were analyzed using one-way ANOVA with Dunnett’s multiple-comparison test, and differences in survival were analyzed using log-rank regression. Fisher’s exact test was performed to examine differences in the numbers of animals with detectable CFU at 12 h.p.i. These statistical analyses were performed using GraphPad Prism version 7 for Macintosh (GraphPad Software, La Jolla, CA). Workspace surfaces were treated with Clorox Healthcare Bleach Germicidal Cleaner to disinfect and prevent spore cross-contamination.

For the murine studies, similar methods were used with the following exceptions. Male and female C57BL/6 mice (*Mus musculus;* Charles River Laboratories) were co-housed, with two to five animals per cage, by treatment group. Instead of clindamycin treatment, mice were provided cefoperazone (0.5 mg/ml; Sigma Aldrich) in drinking water for six days, beginning 8 days prior to inoculation with *C. difficile* (75). Cefoperazone-containing water was exchanged every other day to maintain antibiotic potency. Two days prior to inoculation with *C. difficile*, animals were returned to antibiotic-free sterile drinking water. Spores were prepared as detailed above, and administered as a dose of approximately 1 × 10^5^ spores by oral gavage. Weight loss of ≥ 20% maximum body weight qualified animals as moribund. Differences in CFU counts were analyzed using one-way ANOVA with Dunnett’s multiple-comparison test at each time point, and differences in weight were analyzed using two-way ANOVA with Dunnett’s multiple comparisons test. These statistical analyses were performed using GraphPad Prism version 7 for Macintosh (GraphPad Software, La Jolla, CA).

### Ethics Statement

All animal experimentation was performed under the guidance of veterinarians and trained animal technicians within the Emory University Division of Animal Resources (DAR). Animal experiments were performed with prior approval from the Emory University Institutional Animal Care and Use Committee (IACUC) under protocol #DAR-2001737-052415BA. Animals considered moribund were euthanized by CO_2_ asphyxiation followed by thoracotomy in accordance with the Panel on Euthanasia of the American Veterinary Medical Assocation. The University is in compliance with state and federal Animal Welfare Acts, the standards and policies of the Public Health Service, including documents entitled ‘Guide for the Care and Use of Laboratory Animals’ - National Academy Press, 2011, ‘Public Health Service Policy on Humane Care and Use of Laboratory Animals’ - September 1986, and Public Law 89-544 with subsequent amendments. Emory University is registered with the United States Department of Agriculture (57-R-003) and has filed an Assurance of Compliance statement with the Office of Laboratory Animal Welfare of the National Institutes of Health (A3180-01).

### Germination assays

Spores were purified as described previously, with some modifications (19, 76). *C. difficile* strains were grown on 70:30 sporulation agar plates for 72 h to induce spore formation and allow for vegetative cell lysis. Cells were then scraped from the agar plates, resuspended in sterile water, briefly frozen at −80°C, thawed at 37°C, and left overnight at room temperature. Spore preparations were pelleted at 3200 × g for 20 min, washed in 10 ml of spore stock solution (1x PBS, 1% BSA), pelleted, and resuspended in 1 ml of spore stock solution. The spore suspension was then applied to a 12 ml, 50% sucrose solution and centrifuged at 3200 × g for 20 min. Following centrifugation, the supernatant was decanted and the spore pellet was checked by phase contrast microscopy to verify the elimination of vegetative cells. Sucrose purification was repeated, if necessary, to achieve >95% spore purity. Purified spores were diluted in spore stock solution to a stock concentration of OD_600_ = 3.0. Spores were heat activated for 30 min at 60°C immediately prior to germination assessments. Activated spores were then diluted 1:10 into 800 µl BHIS with either 100 µl of 50 mM taurocholic acid or 100 µl dH_2_O as a negative control, and the OD_600_ was then recorded every 2 min for 20 min. Assays were carried out at room temperature. The percentage decrease in optical density was determined based on the starting OD_600_ for each sample. Assays were performed with spores from three independent spore preparations. Data from the three replicates was averaged and analyzed by a one-way ANOVA for each time point.

### Sporulation assays

Sporulation efficiency was assessed as previously described (77). Briefly, mid-log *C. difficile* cultures at OD_600_ = 0.05 were plated on 70:30 agar and incubated anaerobically at 37°C for 24 hours. Cells were scraped from the plate, resuspended in BHIS, and imaged on a Nikon Eclipse Ci-L microscope with an X100 Ph3 oil-immersion objective. At least 1,000 cells from at least 2 fields of view were assessed per strain and experiment. The percentage of spores was calculated as the number of spores divided by the total number of cells, multiplied by 100. The mean percentage of spores and the standard error of the mean were calculated from three independent experiments and analyzed by two-way ANOVA.

### Accession numbers

*C. difficile* strain 630 (GenBank accession NC_009089.1); *C. difficile* strain R20291 (NC_013316.1). The locus tags for individual genes mentioned in the text are listed in **Table S10**.

## Acknowledgements

The authors would like to thank the members of the McBride lab for useful feedback on this manuscript and Graeme Conn for advice on apparent *K*_d_ calculations. In addition, the authors would like to thank Mike Billingsley, Gregory Tharp, Nirav Patel, and Steven Bosinger of the Yerkes Genomics Core Laboratory for assistance with the next-generation sequencing experiments and data analysis. We thank David Smith and Yi Lasanajak of the Emory Comprehensive Glycomics Core for assistance with the SPR experiments and data analysis. The authors are also grateful to Anice Lowen for the use of her ddPCR machine.

## SUPPORTING INFORMATION-CAPTIONS

**Figure S1. *clnRAB* is transcribed as an operon**. A) The organization of *CD630_16160-CD630_16190* (*clnRAB*). Cultures of strain 630Δ*erm* were grown with or without the addition of 1 µg/ml LL-37, samples collected for RNA and cDNA generated as described in Methods. PCR was performed using genomic DNA (gDNA, positive controls), cDNA templates, or cDNA without reverse transcriptase (RT-, negative controls). Products were generated using primers B) located at the 3’ end of *CD630_16160* (oMC1427) and at the 5’ end of *clnR* (oMC1291), C) within *clnR* (oMC1290) and at the 5’ end of *clnA* (oMC1293), or D) within *clnA* (oMC1292) and at the 5’ end of *clnB* (oMC1295). 25 cycles of PCR were performed and the product visualized on a 0.7% agarose gel. Arrows indicate the expected product size.

**Figure S2. Confirmation of *clnR* and *clnA* mutants, and their complemented strains. A)** PCR products were generated using primers flanking the Targetron *erm::*RAM insertion sites of *clnR* or *clnA*, and genomic DNA from 630Δ*erm* (control)*, clnR* (MC885)*, clnR* Tn:*clnRAB* (MC950)*, clnA* (MC935), or *clnA* Tn:*clnRAB* (MC953) templates. The wild-type PCR product for *clnR* is 696 bp (primers oMC1416/oMC1297), and 558 bp for *clnA* (primers oMC1410/oMC1483). Mutants with intron insertions generate ∼2 kb larger product. Complemented strains yield both the wild-type and insertion products. **B)** Genotypes and phenotypes of mutants and complement strains.

**Figure S3. Alignment of mature cathelicidin sequences.** Cathelicidins from humans (LL-37), mice (mCRAMP) and sheep (SMAP-29) were aligned using the CLUSTAL format alignment (MAFFT, V7.310). Residues corresponding to hydrophobic side chains are labeled above the sequence (Wang *et al*. 2008).

**Figure S4. Alignment of ClnR and GntR-family proteins. A)** Unrooted phylogenetic tree featuring GntR and sub-family representatives: GntR (*B. subtilis;* CAB16042.1), FadR (*T. maritima*; NP_228249.1), HutC (*P. putida*; ADR62377.1), YtrA (*B. subtilis*; KIX83587.1) and ClnR (*C. difficile*; YP_001088118.1) generated using Phylo.io version 1.0k (http://phylo.io/; Robinson et al., 2016 arXiv:1602.04258). **B)** Protein sequences for *C. difficile* ClnR (YP_001088118.1) and *B. subtilis* YtrA (KIX83587.1) were analyzed using MAFFT version 7 (Katoh et al., 2005 Nucl Acids Res). An asterisk below the sequence indicates identical residues, colons indicate similar residues, and dots indicate low similarity.

**Figure S5. Toxin expression in *clnR* and *clnAB* mutant infected hamsters at the time of morbidity.** RNA was extracted from cecal contents of hamsters infected with strain 630Δ*erm,* the *clnR* mutant (MC885), or the *clnAB* mutant (MC935) at the time of morbidity. RNA was used to generate cDNA and analyzed by droplet digital PCR, as described in Methods. Values shown are absolute copies of *tcdA* (A), *tcdB* (B), and *clnR* (C) detected per ng of RNA. Solid lines indicate the median. * indicates adjusted *p* value < 0.05, compared to 630Δ*erm* by one-way ANOVA with Dunnett’s multiple comparisons test.

**Figure S6. Daily fecal CFU counts from mice infected with *cln* mutants.** C57BL/6 mice were inoculated with approximately 1 × 10^5^ spores of 630Δ*erm* (n = 10), *clnR* (MC885; n = 10), *clnAB* (MC935; n = 10), *clnR* Tn::*clnRAB* (MC950; n = 9)), or *clnAB* Tn::*clnRAB* (MC953; n = 9) and CFU in fecal contents were assessed daily, as detailed in Methods. Shown are total *C. difficile* CFU recovered from fecal samples. Dotted line demarcates the lowest limit of detection. Boxes demarcate the interquartile range with median marked by a line. Whiskers extend to the minimum and maximum values. Data were analyzed by one-way ANOVA at each time point, comparing to 630Δ*erm* by Dunnett’s multiple comparison test. * indicates adjusted *p* value < 0.05, ** indicates < 0.002, and *** indicates < 0.0002.

**Figure S7. *clnR* and *clnA* mutant germination and sporulation. A)** Germination assessment for strains 630Δ*erm, clnR* (MC885), and *clnAB* (MC935). Spores were purified as described in methods. Heat-activated spores were added to BHIS for a starting OD_600_ of approximately 0.3. Taurocholic acid (5 mM) was added at T_0_ to the indicated samples, and the OD_600_ of the samples was assessed every two minutes for the duration of the experiment. Ratios of the OD_600_ at each timepoint (T_x_) were plotted against the density observed at T_0_. Three independent biological replicates are shown with error bars indicating the SD. * indicates an adjusted *p* value ≤ 0.05 by one-way ANOVA and Dunnett’s test for multiple comparisons, comparing the mutant strains to 630Δ*erm* (*p* ≤ 0.05 only for *clnR*). **B)** Sporulation frequency. Ethanol-resistant spore formation frequency per total viable CFU of the 630Δ*erm, clnR,* and *clnAB* strains grown on 70:30 sporulation agar with or without 1 µg/ml LL-37 for 24 h. Sporulation frequencies were calculated as described in the Methods. The mean and standard error of the mean are shown for a minimum of three independent experiments. No statistically significant differences were observed by ANOVA assessment.

**Figure S8. LL-37 impacts toxin expression in a ClnRAB-dependent manner.** qRT-PCR analysis of *tcdA* and *tcdB* from strains 630Δ*erm, clnR* (MC885)*, clnAB* (MC935), *clnR* Tn::*clnRAB* (MC950), and *clnAB* Tn::*clnRAB* (MC953) grown in BHIS or BHIS with 2 µg/ml LL-37. Graphs show the mean mRNA expression of **A)** *tcdA* and **B)** *tcdB* relative to expression for 630Δ*erm* in BHIS. Error bars represent the standard error of the mean. Data were analyzed by one-way ANOVA and Dunnett’s multiple comparisons test for comparisons to 630Δ*erm* in the same condition, or by Student’s t-test with Holm-Sidak correction for comparisons of the same strain with/without LL-37. * indicates adjusted *p* value of < 0.05, ** indicates adjusted *P* value of < 0.002. **C)** TcdA western blot for culture lysates from strains grown 24 h in TY medium with or without 2 µg/ml LL-37. Results shown are representative of three independent replicates. Data were analyzed by one-way ANOVA and Dunnett’s multiple comparisons test for comparisons to 630Δ*erm* in the same condition or by Student’s *t* test with Holm-Sidak correction for comparisons of the same strain with/without LL-37. * indicates adjusted *p* value < 0.05 compared to 630Δ*erm* in the same condition, † indicates adjusted *p* value < 0.05 compared to the same strain without LL-37.

**Figure S9. Features of the sequence upstream of *clnR*.** 100 nucleotides upstream of the *clnR* translational start site (TSS in bold) are shown. Underlines indicate putitive −35 and −10 promoter sites. The italicized red region is a tandem repeat sequence that we hypothesize is a ClnR binding site. The construct used in EMSA experiments contained the 84 base pairs upstream of the transcriptional start site, with the final nucleotide indicated in gray.

**Figure S10. Competitive Electrophoretic Mobility Shift Assays (EMSA) of ClnR-dependent genes.** Competitive elecrophoretic mobility shift assays were performed as described in Methods using His-tagged ClnR and fluorescein-labeled DNA encompassing regions upstream of **A**) *iorA,* **B**) *grdE or* **C**) *mtlA*. ClnR was added at 8 µM and specificity was assessed with the addition of unlabeled target DNA or unlabeled P*spo0A* DNA at either 10x or 100x the concentration of labeled target DNA.

**Table S1. Genes differentially expressed in LL-37**.

**Table S2. LL-37 MIC and MBC values for *clnR* and *clnAB* mutants.**

**Table S3. MIC values for *clnR* and *clnAB* mutants in various antimicrobials**.

**Table S4. Genes differentially expressed in a *clnR* mutant in the presence/absence of LL-37.**

**Table S5. Genes regulated by both ClnR and LL-37.**

**Table S6. Relative expression of selected RNA-seq transcripts in *clnR* and *clnAB* mutants.**

**Table S7. Doubling times for 630Δ*erm*, *clnR*, *clnAB*, *clnR* Tn::*clnRAB,* and *clnAB* Tn::*clnRAB in* minimal media supplemented with metabolites, with or without LL-37**

**Table S8. Expression of toxin regulation-associated genes**.

**Table S9. Plasmids and Strains**

**Table S10. Oligonucleotides**

**Table S11. Plasmid construct details.**

**Table S12. Expression of ClnR-dependent genes in the *clnR* mutant complemented with His-ClnR.**

## REFERENCES

1. Lessa FC, Mu Y, Bamberg WM, Beldavs ZG, Dumyati GK, Dunn JR, et al. Burden of *Clostridium difficile* infection in the United States. N Engl J Med. 2015;372(9):825–34.

2. Postma N, Kiers D, Pickkers P. The challenge of *Clostridium difficile* infection: Overview of clinical manifestations, diagnostic tools and therapeutic options. Int J Antimicrob Agents. 2015.

3. Borriello SP. The influence of the normal flora on *Clostridium difficile* colonisation of the gut. Ann Med. 1990;22(1):61–7.

4. Rolfe RD, Helebian S, Finegold SM. Bacterial interference between *Clostridium difficile* and normal fecal flora. J Infect Dis. 1981;143(3):470–5.

5. Peschel A, Sahl HG. The co-evolution of host cationic antimicrobial peptides and microbial resistance. Nat Rev Microbiol. 2006;4(7):529–36.

6. Sorensen O, Arnljots K, Cowland JB, Bainton DF, Borregaard N. The human antibacterial cathelicidin, hCAP-18, is synthesized in myelocytes and metamyelocytes and localized to specific granules in neutrophils. Blood. 1997;90(7):2796–803.

7. Chen CI, Schaller-Bals S, Paul KP, Wahn U, Bals R. Beta-defensins and LL-37 in bronchoalveolar lavage fluid of patients with cystic fibrosis. J Cyst Fibros. 2004;3(1):45–50.

8. Raqib R, Sarker P, Mily A, Alam NH, Arifuzzaman AS, Rekha RS, et al. Efficacy of sodium butyrate adjunct therapy in shigellosis: a randomized, double-blind, placebo-controlled clinical trial. BMC Infect Dis. 2012;12:111.

9. Dürr UHN, Sudheendra US, Ramamoorthy A. LL-37, the only human member of the cathelicidin family of antimicrobial peptides. BBA - Biomembranes. 2006;1758(9):1408–25.

10. Cowardin CA, Petri WA, Jr. Host recognition of *Clostridium difficile* and the innate immune response. Anaerobe. 2014;30:205–9.

11. Kelly CP, Kyne L. The host immune response to *Clostridium difficile*. J Med Microbiol. 2011;60(Pt 8):1070–9.

12. McQuade R, Roxas B, Viswanathan VK, Vedantam G. *Clostridium difficile* clinical isolates exhibit variable susceptibility and proteome alterations upon exposure to mammalian cationic antimicrobial peptides. Anaerobe. 2012;18(6):614–20.

13. Kazamias MT, Sperry JF. Enhanced fermentation of mannitol and release of cytotoxin by *Clostridium difficile* in alkaline culture media. Appl Environ Microbiol. 1995;61(6):2425–7.

14. Janoir C, Deneve C, Bouttier S, Barbut F, Hoys S, Caleechum L, et al. Adaptive strategies and pathogenesis of *Clostridium difficile* from *in vivo* transcriptomics. Infect Immun. 2013;81(10):3757–69.

15. Ferreyra JA, Wu KJ, Hryckowian AJ, Bouley DM, Weimer BC, Sonnenburg JL. Gut microbiota-produced succinate promotes *C. difficile* infection after antibiotic treatment or motility disturbance. Cell host & microbe. 2014;16(6):770–7.

16. Ho TD, Ellermeier CD. PrsW is required for colonization, resistance to antimicrobial peptides, and expression of extracytoplasmic function sigma factors in *Clostridium difficile*. Infect Immun. 2011;79(8):3229–38.

17. Theriot CM, Koenigsknecht MJ, Carlson PE, Jr., Hatton GE, Nelson AM, Li B, et al. Antibiotic-induced shifts in the mouse gut microbiome and metabolome increase susceptibility to *Clostridium difficile* infection. Nat Commun. 2014;5:3114.

18. Dineen SS, McBride SM, Sonenshein AL. Integration of metabolism and virulence by *Clostridium difficile* CodY. J Bacteriol. 2010;192(20):5350–62.

19. Nawrocki KL, Edwards AN, Daou N, Bouillaut L, McBride SM. CodY-dependent regulation of sporulation in *Clostridium difficile*. J Bacteriol. 2016;198(15):2113–30.

20. Edwards AN, Nawrocki KL, McBride SM. Conserved oligopeptide permeases modulate sporulation initiation in *Clostridium difficile*. Infect Immun. 2014;82(10):4276–91.

21. Woods EC, McBride SM. Regulation of antimicrobial resistance by extracytoplasmic function (ECF) sigma factors. Microbes Infect. 2017;19(4-5):238–48.

22. Woods EC, Nawrocki KL, Suarez JM, McBride SM. The *Clostridium difficile* Dlt pathway is controlled by the ECF sigma factor, SigmaV, in response to lysozyme. Infect Immun. 2016.

23. Nawrocki KL, Wetzel D, Jones JB, Woods EC, McBride SM. Ethanolamine is a valuable nutrient source that impacts *Clostridium difficile* pathogenesis. Environ Microbiol. 2018;20(4):1419–35.

24. Utaida S, Dunman PM, Macapagal D, Murphy E, Projan SJ, Singh VK, et al. Genome-wide transcriptional profiling of the response of *Staphylococcus aureus* to cell-wall-active antibiotics reveals a cell-wall-stress stimulon. Microbiology. 2003;149(Pt 10):2719–32.

25. Poole K. Bacterial stress responses as determinants of antimicrobial resistance. J Antimicrob Chemother. 2012;67(9):2069–89.

26. Durr UH, Sudheendra US, Ramamoorthy A. LL-37, the only human member of the cathelicidin family of antimicrobial peptides. Biochim Biophys Acta. 2006;1758(9):1408–25.

27. Skerlavaj B, Benincasa M, Risso A, Zanetti M, Gennaro R. SMAP-29: a potent antibacterial and antifungal peptide from sheep leukocytes. FEBS Lett. 1999;463(1-2):58–62.

28. Wang G. Structures of human host defense cathelicidin LL-37 and its smallest antimicrobial peptide KR-12 in lipid micelles. J Biol Chem. 2008;283(47):32637–43.

29. Hoskisson PA, S. Rigali Chapter 1: Variation in form and function the helix-turn-helix regulators of the GntR superfamily. Adv Appl Microbiol. 2009;69:1–22.

30. Rigali S, Derouaux A, Giannotta F, Dusart J. Subdivision of the helix-turn-helix GntR family of bacterial regulators in the FadR, HutC, MocR, and YtrA subfamilies. J Biol Chem. 2002;277(15):12507–15.

31. Bouillaut L, Self WT, Sonenshein AL. Proline-dependent regulation of *Clostridium difficile* Stickland metabolism. J Bacteriol. 2013;195(4):844–54.

32. Dubois T, Dancer-Thibonnier M, Monot M, Hamiot A, Bouillaut L, Soutourina O, et al. Control of *Clostridium difficile* physiopathology in response to cysteine availability. Infect Immun. 2016;84(8):2389–405.

33. Karlsson S, Burman LG, Akerlund T. Suppression of toxin production in *Clostridium difficile* VPI 10463 by amino acids. Microbiology. 1999;145 (Pt 7):1683–93.

34. Keel MK, Songer JG. The comparative pathology of *Clostridium difficile*-associated disease. Vet Pathol. 2006;43(3):225–40.

35. Chang TW, Bartlett JG, Gorbach SL, Onderdonk AB. Clindamycin-induced enterocolitis in hamsters as a model of pseudomembranous colitis in patients. Infect Immun. 1978;20(2):526–9.

36. Bartlett JG, Onderdonk AB, Cisneros RL, Kasper DL. Clindamycin-associated colitis due to a toxin-producing species of *Clostridium* in hamsters. J Infect Dis. 1977;136(5):701–5.

37. Larson HE, Price AB, Honour P, Borriello SP. Clostridium difficile and the aetiology of pseudomembranous colitis. Lancet. 1978;1(8073):1063–6.

38. Chen X, Katchar K, Goldsmith JD, Nanthakumar N, Cheknis A, Gerding DN, et al. A mouse model of *Clostridium difficile*-associated disease. Gastroenterology. 2008;135(6):1984–92.

39. Best EL, Freeman J, Wilcox MH. Models for the study of *Clostridium difficile* infection. Gut Microbes. 2012;3(2):145–67.

40. Martin-Verstraete I, Peltier J, Dupuy B. The regulatory networks that control *Clostridium difficile* toxin synthesis. Toxins (Basel). 2016;8(5).

41. Gellatly SL, Needham B, Madera L, Trent MS, Hancock RE. The *Pseudomonas aeruginosa* PhoP-PhoQ two-component regulatory system is induced upon interaction with epithelial cells and controls cytotoxicity and inflammation. Infect Immun. 2012;80(9):3122–31.

42. Froehlich BJ, Bates C, Scott JR. *Streptococcus pyogenes* CovRS mediates growth in iron starvation and in the presence of the human cationic antimicrobial peptide LL-37. J Bacteriol. 2009;191(2):673–7.

43. Nizet V, Ohtake T, Lauth X, Trowbridge J, Rudisill J, Dorschner RA, et al. Innate antimicrobial peptide protects the skin from invasive bacterial infection. Nature. 2001;414(6862):454–7.

44. Velarde JJ, Ashbaugh M, Wessels MR. The human antimicrobial peptide LL-37 binds directly to CsrS, a sensor histidine kinase of group A *Streptococcus*, to activate expression of virulence factors. J Biol Chem. 2014;289(52):36315–24.

45. Tran-Winkler HJ, Love JF, Gryllos I, Wessels MR. Signal transduction through CsrRS confers an invasive phenotype in group A *Streptococcus*. PLoS Pathog. 2011;7(10):e1002361.

46. Majchrzykiewicz JA, Kuipers OP, Bijlsma JJ. Generic and specific adaptive responses of *Streptococcus pneumoniae* to challenge with three distinct antimicrobial peptides, bacitracin, LL-37, and nisin. Antimicrob Agents Chemother. 2010;54(1):440–51.

47. Strempel N, Neidig A, Nusser M, Geffers R, Vieillard J, Lesouhaitier O, et al. Human host defense peptide LL-37 stimulates virulence factor production and adaptive resistance in *Pseudomonas aeruginosa*. PLoS One. 2013;8(12):e82240.

48. Liu W, Dong SL, Xu F, Wang XQ, Withers TR, Yu HD, et al. Effect of intracellular expression of antimicrobial peptide LL-37 on growth of *Escherichia coli* strain TOP10 under aerobic and anaerobic conditions. Antimicrob Agents Chemother. 2013;57(10):4707–16.

49. Sonenshein AL. CodY, a global regulator of stationary phase and virulence in Gram-positive bacteria. Curr Opin Microbiol. 2005;8(2):203–7.

50. Dineen SS, Villapakkam AC, Nordman JT, Sonenshein AL. Repression of *Clostridium difficile* toxin gene expression by CodY. Mol Microbiol. 2007;66(1):206–19.

51. Dupuy B, Sonenshein AL. Regulated transcription of *Clostridium difficile* toxin genes. Mol Microbiol. 1998;27(1):107–20.

52. Antunes A, Martin-Verstraete I, Dupuy B. CcpA-mediated repression of *Clostridium difficile* toxin gene expression. Mol Microbiol. 2011;79(4):882–99.

53. Antunes A, Camiade E, Monot M, Courtois E, Barbut F, Sernova NV, et al. Global transcriptional control by glucose and carbon regulator CcpA in *Clostridium difficile*. Nucleic Acids Res. 2012;40(21):10701–18.

54. Bouillaut L, Dubois T, Sonenshein AL, Dupuy B. Integration of metabolism and virulence in *Clostridium difficile*. Res Microbiol. 2015;166(4):375–83.

55. Karlsson S, Burman LG, Akerlund T. Induction of toxins in *Clostridium difficile* is associated with dramatic changes of its metabolism. Microbiology. 2008;154(Pt 11):3430–6.

56. Pitts AC, Tuck LR, Faulds-Pain A, Lewis RJ, Marles-Wright J. Structural insight into the *Clostridium difficile* ethanolamine utilisation microcompartment. PLoS One. 2012;7(10):e48360.

57. Kopke M, Straub M, Durre P. *Clostridium difficile* is an autotrophic bacterial pathogen. PLoS One. 2013;8(4):e62157.

58. Luria SE, Burrous JW. Hybridization between *Escherichia coli* and *Shigella*. J Bacteriol. 1957;74(4):461–76.

59. Putnam EE, Nock AM, Lawley TD, Shen A. SpoIVA and SipL are *Clostridium difficile* spore morphogenetic proteins. J Bacteriol. 2013;195(6):1214–25.

60. Edwards AN, Suarez JM, McBride SM. Culturing and maintaining *Clostridium difficile* in an anaerobic environment. J Vis Exp. 2013(79):e50787.

61. Bouillaut L, McBride SM, Sorg JA. Genetic manipulation of *Clostridium difficile*. Curr Protoc Microbiol. 2011;Chapter 9:Unit 9A 2.

62. Sorg JA, Dineen SS. Laboratory maintenance of *Clostridium difficile*. Curr Protoc Microbiol. 2009;Chapter 9:Unit9A 1.

63. Suarez JM, Edwards AN, McBride SM. The *Clostridium difficile cpr* locus is regulated by a noncontiguous two-component system in response to type A and B lantibiotics. J Bacteriol. 2013;195(11):2621–31.

64. Dobin A, Davis CA, Schlesinger F, Drenkow J, Zaleski C, Jha S, et al. STAR: ultrafast universal RNA-seq aligner. Bioinformatics. 2013;29(1):15–21.

65. Anders S, Pyl PT, Huber W. HTSeq--a Python framework to work with high-throughput sequencing data. Bioinformatics. 2015;31(2):166–9.

66. Love MI, Huber W, Anders S. Moderated estimation of fold change and dispersion for RNA-seq data with DESeq2. Genome Biol. 2014;15(12):550.

67. Tatusov RL, Koonin Ev Fau-Lipman DJ, Lipman DJ. A genomic perspective on protein families. Science. 1997(0036-8075 (Print)).

68. Schmittgen TD, Livak KJ. Analyzing real-time PCR data by the comparative C(T) method. Nat Protoc. 2008;3(6):1101–8.

69. Liu R, Suarez JM, Weisblum B, Gellman SH, McBride SM. Synthetic polymers active against *Clostridium difficile* vegetative cell growth and spore outgrowth. J Am Chem Soc. 2014;136(41):14498–504.

70. Edwards AN, Tamayo R, McBride SM. A novel regulator controls *Clostridium difficile* sporulation, motility and toxin production. Mol Microbiol. 2016.

71. Mercante J, Suzuki K, Cheng X, Babitzke P, Romeo T. Comprehensive alanine-scanning mutagenesis of *Escherichia coli* CsrA defines two subdomains of critical functional importance. J Biol Chem. 2006;281(42):31832–42.

72. Cartman ST, Kelly ML, Heeg D, Heap JT, Minton NP. Precise manipulation of the *Clostridium difficile* chromosome reveals a lack of association between the *tcdC* genotype and toxin production. Appl Environ Microbiol. 2012;78(13):4683–90.

73. Healthcare G. Biacore Sensor Surface Handbook. 2007.

74. Wang Y, Agerberth B, Lothgren A, Almstedt A, Johansson J. Apolipoprotein A-I binds and inhibits the human antibacterial/cytotoxic peptide LL-37. J Biol Chem. 1998;273(50):33115–8.

75. Theriot CM, Koumpouras CC, Carlson PE, Bergin, II, Aronoff DM, Young VB. Cefoperazone-treated mice as an experimental platform to assess differential virulence of *Clostridium difficile* strains. Gut Microbes. 2011;2(6):326–34.

76. Sorg JA, Sonenshein AL. Inhibiting the initiation of *Clostridium difficile* spore germination using analogs of chenodeoxycholic acid, a bile acid. J Bacteriol. 2010;192(19):4983–90.

77. Edwards AN, McBride SM. Isolating and Purifying *Clostridium difficile* Spores. Methods Mol Biol. 2016;1476:117–28.

